# Classification models distinguish functional and trafficking effects of KCNQ1 variants to enhance variant interpretation

**DOI:** 10.1101/2025.10.31.685955

**Authors:** Ana C. Chang-Gonzalez, Eric W. Bell, Carlos G. Vanoye, Eduardo Guadarrama, Reshma R. Desai, Jean-Marc DeKeyser, Kathryn R. Butcher, Thomas Scott, Charles R. Sanders, Alfred L. George, Kaitlyn V. Ledwitch, Jens Meiler

## Abstract

Missense mutations compromise protein fitness by altering stability and function, which can lead to various clinical disease states. The potassium ion channel KCNQ1 underlies the majority of congenital long QT syndrome (LQTS) cases, one of the most common genetic arrhythmia syndromes. During genetic testing for LQTS, variants of uncertain significance (VUS) confound diagnosis and clinical management. KCNQ1 protein fitness metrics enable mechanistic classification of variants, directly informing the molecular basis for dysfunction and providing clinical interpretation of variants linked to LQTS and other channelopathies. We developed structure-aware random forest classifier models to predict seven metrics of KCNQ1 fitness, four functional electrophysiology measurements (peak current density, voltage-dependence, gating kinetics), and three trafficking values measured by flow cytometry. Our trained models outperformed AlphaMissense in predicting protein fitness, enhancing interpretation of ClinVar VUS and variants classified as ambiguous by AlphaMissense. We demonstrate the classifiers distinguish benign and pathogenic variants from ClinVar and gnomAD and identify systematic patterns of dysfunction and mistrafficking along the functionally critical S4 helix. Our method advances variant effect prediction with a mechanistic classifier that reliably links missense mutations in KCNQ1 to their specific disease-causing mechanisms. As a resource for precision medicine approaches for LQTS or other KCNQ1 channelopathies, we provide the predictions and scores for all KCNQ1 missense variants across the structured region of the protein.

## Introduction

Computational variant effect predictors (VEPs) are increasingly relied upon to assess the clinical impact of missense mutations. Recent structure-aware VEPs exhibit better performance over older methods based solely on multiple sequence alignments (MSAs) in distinguishing pathogenic variants [1]. However, VEPs typically assign a general pathogenicity score based on patterns observed in large datasets, without predicting the specific functional or mechanistic outcomes (e.g., changes in protein stability, function, or cellular trafficking) that lead to disease. This lack of mechanistic insight limits their utility in clinical settings where understanding why a variant is pathogenic is necessary for accurate diagnosis and treatment.

Benchmarks confirm this limitation, as published VEPs show variable performance across different protein tasks [2-5]. For instance, AlphaMissense (AM) scores correlated well with the *in vitro* ion current of the CFTR gene but failed to correlate with CFTR trafficking, and only modestly reflected clinical measures of disease severity [6]. The primary reason VEPs lack mechanistic training is the paucity of experimental data. Typically, experimental assays evaluate a few hundred variants per protein target, often concentrating on specific domains or variant classes. This is more acute for membrane proteins, largely because membrane proteins function in a compositionally and biophysically complex lipid environment, making experimental studies challenging and data more limited [7]. Consequently, most VEPs are trained and benchmarked on datasets that overrepresent soluble proteins, raising concerns about their generalizability and suitability for providing mechanistic interpretations in the context of membrane proteins. Currently, only a handful of tools have been developed specifically for membrane proteins [8-10]. The most recent, MutDPAL, aims to improve prediction accuracy by integrating transmembrane environment features for multi-label disease classification [9].

Among clinically-relevant membrane proteins is the human voltage-gated K^+^ ion channel K_V_7.1 (KCNQ1), a homotetramer where each 676-residue monomer contains a voltage-sensing domain (VSD) and a pore domain (PD) [11]. K_V_7.1 complexes with the accessory subunit KCNE1 to generate the slow delayed rectifier current (*I*_Ks_) essential for normal myocardial repolarization [12, 13]. KCNQ1 mutations are associated with the majority of cases of the genetic cardiac arrhythmia, congenital long QT syndrome (LQTS) [14-16]. Patients with KCNQ1-associated LQTS have structurally normal hearts, yet external triggers can cause life-threatening cardiac events [15]. While genetic testing helps in the diagnosis of LQTS, rare variants and clinically silent mutations often complicate variant interpretation [17].

Pathogenic KCNQ1 variants cause LQTS through diverse mechanisms, the most common being channel mistrafficking [18-20]. In previous work, we grouped KCNQ1 variants into six mechanistic classes based on functional measurements from electrophysiological recordings, stability measurements, and channel expression/trafficking from flow cytometry-based trafficking assay [19, 20]. We found that variants in the VSD and PD associated with LQTS sometimes traffic normally but impair channel function by reducing current density, shifting voltage dependence, or altering gating kinetics. In other cases, disease variants exhibited reduced trafficking to the membrane surface as the cause of reduced channel function. Therefore, these classes capture distinct molecular modes of KCNQ1 dysfunction, defined by characteristic patterns across both functional and trafficking measurements. Because variants in each class span KCNQ1 structural domains and differ in the type of amino acid substitution. Therefore, establishing rule-based criteria from annotated variants to classify new uncharacterized variants into mechanistic categories is not a trivial task.

Here, we present a machine learning framework to predict variant effects on KCNQ1 function and trafficking to facilitate the interpretation of VUS. Using curated features that capture the structural and evolutionary impact of each variant, we trained random forest (RF) classifiers to predict binary outcomes for each function and trafficking measurement. The labels we used for variant classification were collected by measuring properties of the mutant channel relative to WT protein. We report the functional properties of nearly 100 previously uncharacterized variants, half of which are VUS. We demonstrate the utility of our model predictions by accurately predicting the functional impact of ClinVar VUS, and interpreting the mechanism of variants classified as “ambiguous” by AM [21] or “uncertain” by electrophysiological measurements. We developed a global score of dysfunction and a global score of mistrafficking from the independent protein fitness metric predictions that discriminates annotated benign and pathogenic variants from ClinVar [22] and gnomAD [23]. Our classifiers use features shared across membrane proteins, making the framework transferable to other membrane proteins. All scripts, training data, model weights, and predictions are made publicly available.

## Results

### Random forest classifier models predict variant functional and trafficking outcomes

The model training set consisted of KCNQ1 variants with functional and trafficking measurements from our previous studies [19, 20, 24, 25] and additional variants we tested for this work (see *Methods*, VUS and PD sets). We strategically selected KCNQ1 variants of uncertain significance (VUS) in ClinVar and variants in the pore domain, a region underrepresented in our previous set [24], to add to the model. In total, we curated 201 variants with electrophysiological data and 145 variants with trafficking measurements (**Fig. 1A**).

**Figure 1.**
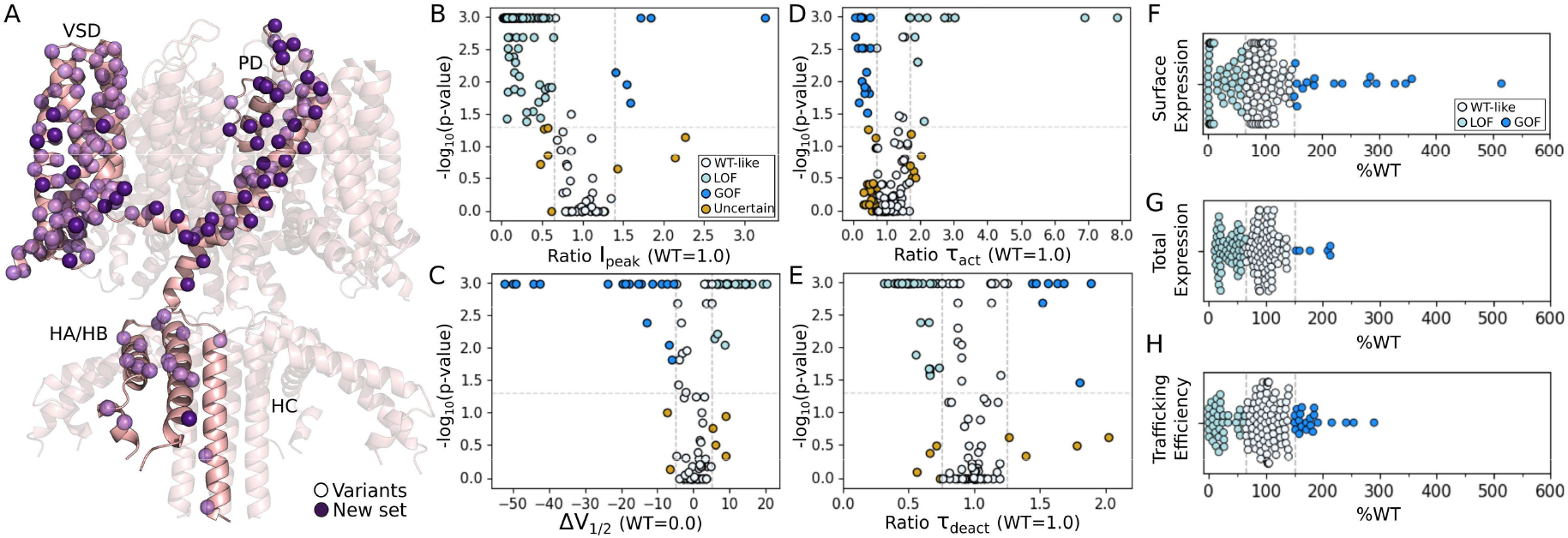
Training data. (**A**) Structural model of KCNQ1 based on PDB 8SIK[37] used to generate biophysical features. Variants used for training and cross-validation studies are mapped onto the structure. Spheres represent variant Cα atoms. Variants functionally characterized for this study are colored dark purple. VSD=voltage-sensing domain. PD=pore domain. HA, HB, HC=intracellular helical domains. S4 set variants are not shown in the mapping. (**B-E**) Functional measurements from electrophysiological recordings. (**F-H**) Trafficking measurements. Function and trafficking measurements were scaled to wild-type (WT). Dashed lines denote cutoff thresholds for classification (**Table 1**) and colors correspond to assigned class labels (**Table 2**) as follows: cyan=loss-of-function (LOF), white=WT-like normal, blue=gain-of-function (GOF), and orange=uncertain. Variants with “Not determined” measurements are not shown. This figure details the variants included in training the RF classifiers.

**Figures 1B-H** plot function and trafficking measurements and respective variant class label assignments. The four electrophysiological measurements used to characterize KCNQ1 function were: peak current density (I_peak_), the midpoint of activation voltage dependence (V_1/2_), and time constants for activation (τ_act_) and deactivation (τ_deact_). The three trafficking measures were: cell surface expression, total expression, and trafficking efficiency. Together, these seven values are referred to as protein fitness metrics. Each variant was assigned a label corresponding to the measurement classification, being either within the range of the WT variant (normal=0) or outside of that range (dysfunctional=1). Labels were assigned using threshold cutoff values and statistical significance following guidance from our previous work [19, 24] (**Table 1**).

**Table 1.** Threshold cutoff values used to classify variants. Collected measurements were relative to WT channel such that 1.0 for I_peak_, τ_act_, τ_deact,_ 0.0 mV for V_1/2_, and 100% for surface and total expression and trafficking efficiency indicate that the variant measurement is equivalent to that of the WT channel, reflecting normal function.

| Measurement | LOF (relative to WT) | GOF (relative to WT) |
| --- | --- | --- |
| $I_{\text{peak}}$ | < 0.65 | > 1.40 |
| $V_{1/2}$ | > 5.0 mV | < -5.0 mV |
| $\tau_{\text{act}}$ | > 1.70 | < 0.70 |
| $\tau_{\text{deact}}$ | < 0.75 | > 1.25 |
| Surface expression | < 65% | > 150% |
| Total expression | < 65% | > 150% |
| Trafficking efficiency | < 65% | > 150% |

**Table 2.** Number of variants per classification used in this work for model development. Total number of variants is 201 for functional metrics and 145 for trafficking metrics. Variants were compiled from our previous studies [20, 24, 25, 36] in addition to new characterized variants. S4 set variants are not included in these counts.

| Measurement | Normal | LOF | GOF | Not determined | Uncertain |
| --- | --- | --- | --- | --- | --- |
| $I_{\text{peak}}$ | 46 | 141 | 6 | 0 | 8 |
| $V_{1/2}$ | 62 | 24 | 18 | 91 | 6 |
| $\tau_{\text{act}}$ | 88 | 14 | 23 | 50 | 26 |
| $\tau_{\text{deact}}$ | 72 | 33 | 8 | 80 | 8 |
| Surface expression | 75 | 53 | 17 | - | - |
| Total expression | 85 | 54 | 6 | - | - |
| Trafficking efficiency | 88 | 36 | 21 | - | - |

We trained individual RF classifiers using curated features from our published KCNQ1 function predictor [24] to predict the seven protein fitness metrics. The features include 11 biophysical values that capture both the physicochemical nature of the amino acid substitution (i.e., changes in hydrogen-bonding potential, van der Waals volume, hydrophobicity, and polarizability) and the structural context of the variant (i.e., distance from the pore axis, residue burial, and local effects), and one evolutionary feature. We refer to this feature set as “BioEvo.” The RF classifiers achieved median Matthews correlation coefficient (MCC) values above 0.5 (**Fig. 2A**, *red* distributions), low Brier scores (**Fig. 2B**, *red* distributions), high area under the receiver operating characteristic curve (AUROC), and high area under the precision-recall curve (AUPRC) values (**Figs. S1B** and **S1C**). Using individual RF classifiers to predict each protein fitness metric performed better than using a multi-label RF model (data not shown), reflecting the independent nature of the phenotypic outcomes that lead to distinct mechanisms of dysfunction [19, 20]. This study resulted in seven trained RF classifiers using the BioEvo features, one to predict each protein fitness metric.

**Figure 2.**
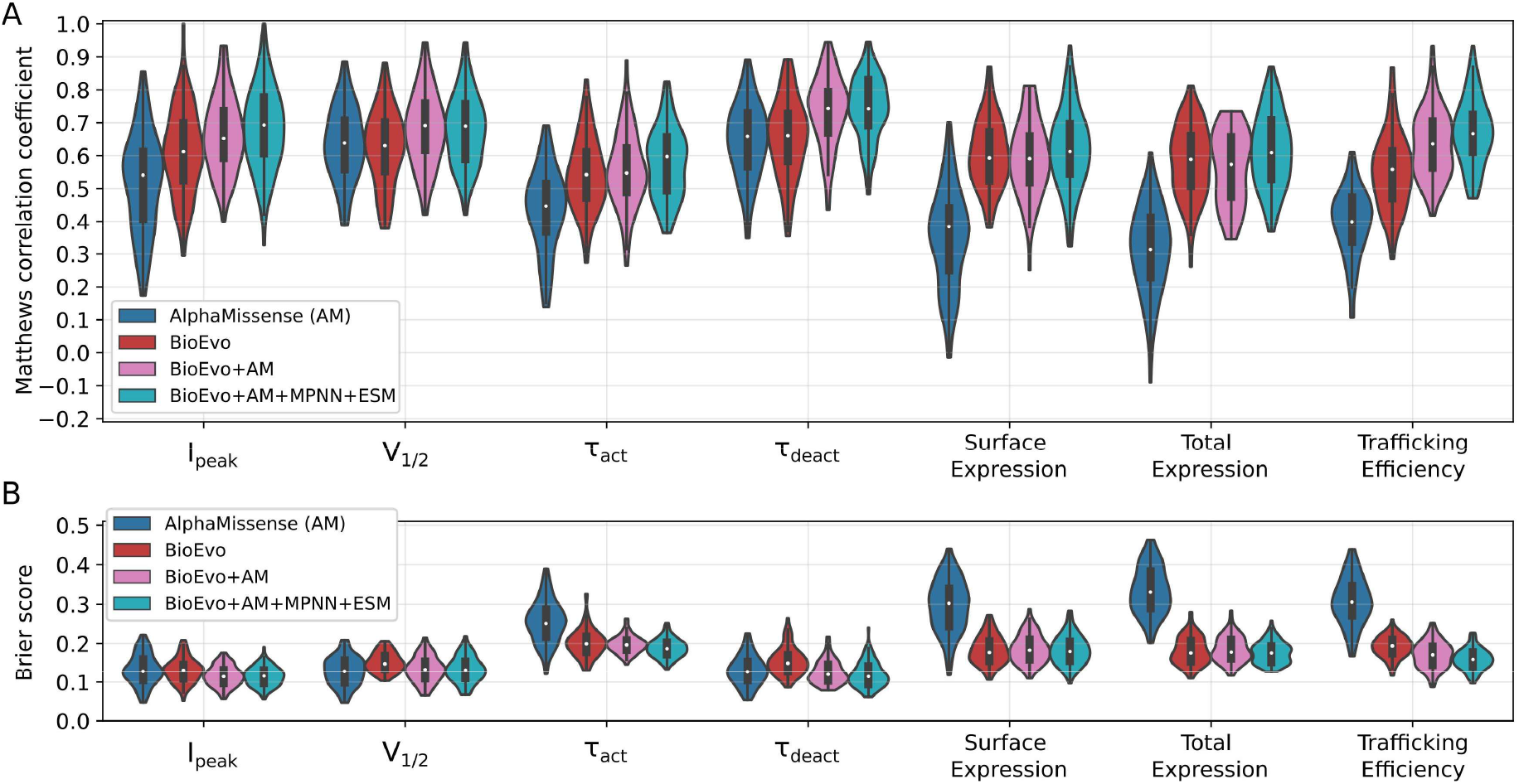
Cross-validation metrics using different feature sets. (**A**) Matthews correlation coefficient (MCC) and (**B**) Brier score to evaluate binary classification model performance. The “AlphaMissense” set uses the AlphaMissense (AM) pathogenicity score [21] directly to classify variants. “BioEvo” are features developed in our previous work [24]. “BioEvo+AM” combine BioEvo and AM, and “BioEvo+AM+MPNN+ESM” combine those plus amino acid likelihood probabilities from ProteinMPNN [29] and ESM [31]. Each distribution contains 100 calculations from the validation sets of 20 repeats of 5-fold cross-validation. The same variants were included in the validation set for I_peak_ across the four model feature sets, making the validation metrics comparable (see **Fig. S1A**). The same applied to V_1/2_ and all other predicted measurements. Variants marked as “uncertain” or “not measured” were excluded from the validation metrics. Cross-validation metrics support the utility of BioEvo+AM+MPNN+ESM features in predicting KCNQ1 function and trafficking, measures that reflect protein fitness.

### AlphaMissense pathogenicity score as an additional feature improves variant effect prediction

We used the pathogenicity scores from AlphaMissense (AM) [21] to test the hypothesis that adding the output of a general pathogenicity predictor as an additional feature to the RF classifiers improves function and trafficking predictions. AM performs well across diverse sets of benchmark proteins [26] and has demonstrated some utility in predicting KCNQ1 variant dysfunction [27, 28]. The AM pathogenicity score is a continuous number from 0 to 1, where 0 indicates that the missense variant is likely benign and 1 that it is likely pathogenic. This score is fine-tuned based on the frequency of missense variants in the human or primate populations, where variants unobserved in the human population are classified as pathogenic [21]. Because population frequency data is not represented in the BioEvo features, the AM pathogenicity score adds orthogonal information to our RF models.

To test AM as a standalone predictor of the seven protein fitness metrics, we used its pathogenicity score as the predicted value for each metric and the published benign/pathogenic classification labels to indicate WT-like or impaired behavior. Because AM reports only one predicted metric (pathogenicity), each variant was assigned the same classification across all metrics. Validation metrics were computed on the same validation sets as the BioEvo model (**Figs. 2** and **S1**, *BioEvo* versus *AlphaMissense*). We calculated the difference in MCC between AM and BioEvo predictions on each validation set to directly compare performance per set. For five out of the seven predicted metrics, the median MCC using the direct AM classifications were 0.1 to 0.2 lower than using the BioEvo RF models (**Fig. 2A**, *blue* versus *red*; **Fig. S1A**, *blue*). For V_1/2_ and τ_deact_, median MCC of the predicted metrics were comparable. MCC values varied more across validation folds using AM compared to BioEvo predictions, especially for trafficking-related metrics (**Fig. 2A**, *blue* versus *red*), where the AM MCC drops below 0, indicating predictions were worse than randomness. This trend is reflected in the comparatively higher Brier scores for trafficking predictions (**Fig. 2B**, *blue* versus *red*). These results indicate improved performance using BioEvo RF models over AM pathogenicity predictions to classify variant function and trafficking.

While BioEvo RF models performed better when evaluated by MCC and Brier scores, direct use of AM resulted in higher AUROC and AUPRC values (**Figs. S1B** and **S1C**). This difference in performance suggests that the two models capture unique and complementary patterns in the data. Consequently, an approach combining the BioEvo features with the prediction from AM could be employed to create a more robust predictor. We retrained and validated the RF classifiers using 13 features, the 12 BioEvo plus AM pathogenicity score (BioEvo+AM). For all predicted metrics, the BioEvo+AM median MCC were higher than either AM or BioEvo individually (**Fig. 2A**, *pink* versus *blue* and *red*). Compared to using AM directly, the BioEvo+AM classifiers consistently improved MCC (**Fig. S1A**, *orange*). With a few exceptions, the BioEvo+AM classifiers increased AUROC and AUPRC over BioEvo features alone (**Figs. S1B** and **S1C**). Together, these data indicate that RF classifiers trained on protein-specific biophysical and evolutionary features plus general pathogenicity predictions improve predictions of KCNQ1 protein fitness metrics beyond either approach alone.

### Incorporating amino acid probabilities from ProteinMPNN and ESM-2 modestly improves prediction of variant effects on KCNQ1 function and trafficking

Building on the improvement from the BioEvo+AM features, we tested whether adding features from protein structure prediction models leads to greater performance gains. For this, we used the protein message passing neural network (ProteinMPNN) and Evolutionary Scale Modeling (ESM-2) amino acid likelihood probabilities. ProteinMPNN is a structure-based deep learning model that takes a protein backbone as input and assigns a position-specific probability that a given amino acid will be stable and structurally comparable as the WT residue at that site [29]. ESM-2 is a sequence-based protein language model that uses evolutionary context learned from a vast database of protein sequences to predict the probability of an amino acid at a given position [30, 31]. We retrained the RF classifiers including the per-variant probabilities from ProteinMPNN and ESM-2 models (BioEvo+AM+MPNN+ESM).

We compared validation metrics between the trained models and across predicted measurements to identify the optimal feature set for the seven classification tasks. For the RF classifiers trained on the expanded 15 BioEvo+AM+MPNN+ESM feature set, median MCC either increased or was comparable to that of the classifiers trained on the other feature sets (**Fig. 2A**, *green*). Compared to using AM directly, the BioEvo+AM+MPNN+ESM classifiers consistently improved MCC (**Fig. S1A**, *green*). Brier scores were comparable for all the models that included BioEvo features (**Fig. 2A**, *red, pink*, and *green*). Using BioEvo+AM+MPNN+ESM features also achieved higher or similar AUROC and AUPRC (**Figs. S1B** and **S1C**) compared to using BioEvo+AM features. RF classifiers trained on protein-specific biophysical and evolutionary features, pathogenicity predictions, and amino acid probabilities from structure prediction models improved prediction performance and overall provided a robust feature set suitable for predicting variant effects on KCNQ1 function and trafficking. The variant predictions reported in the rest of this paper are from RF classifiers trained using the BioEvo+AM+MPNN+ESM features.

### RF classifiers discern functional properties of ClinVar variants of uncertain significance

We applied our published Q1VarPredBio model to predict functional outcomes for KCNQ1 missense variants annotated in ClinVar as VUS. From these, we selected a balanced subset of 23 variants for functional evaluation, using predictions which indicated that half of the selected variants should have WT-like function. Excluding these 23 VUS, we re-trained the RF classifiers to avoid data leakage and minimize bias toward the test set. For the classifier of each metric, we performed 100 shuffled splits with stratified sampling, splitting the remaining variants into 80% training and 20% validation sets. MCC values were computed on the validation set for each split. We used the RF model with the highest MCC to predict protein fitness for these VUS (**Fig. 3A**). Trafficking predictions are included for reference, but trafficking data were not collected for this subset.

**Figure 3.**
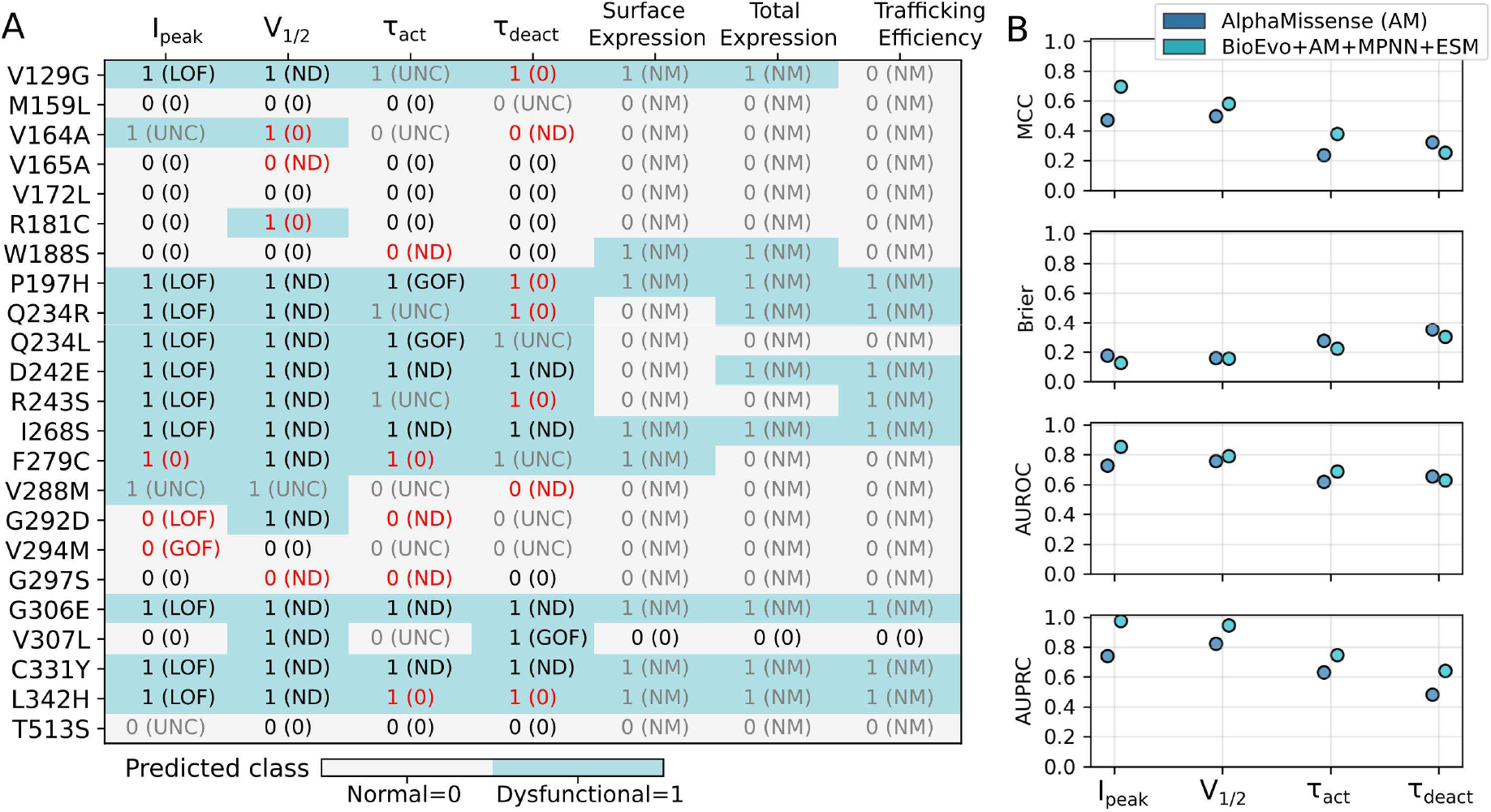
Model predictions on a subset of ClinVar variants of uncertain significance (VUS). (**A**) Heatmap where dysfunctional predictions are marked by blue cells. Text annotations are the predicted classes, and the true labels are in parentheses. LOF, GOF, and ND were all considered dysfunctional. Red text indicates the prediction does not match the true label. Gray text indicates that the corresponding metric was either not measured (“NM”) or true label was uncertain (“UNC”). ClinVar was accessed before 2024 to collect the list of VUS. As of August 7, 2025, D242E and R243S were annotated in ClinVar as either likely pathogenic or pathogenic. (**B**) Validation metrics on the variants in panel A using the AM pathogenicity scores or our RF classifiers (BioEvo+AM+MPNN+ESM). “UNC” and “NM” measurements were excluded from metric calculations. Performance metrics for trafficking measurements are not shown as most of these were not measured (marked “NM” in panel A). RF classifiers improve function predictions for ClinVar VUS over using AM pathogenicity score alone.

The RF classifiers trained on BioEvo+AM+MPNN+ESM features outperform classifications of the 23 VUS using AM predictions directly (**Fig. 3B**) and predictions using Q1VarPredBio (**Fig. S2**). This agrees with our findings from the cross-validation performance on the full set of variants (**Figs. 2** and **S1**). Since the test set variants were explicitly excluded from model training, this demonstrates that the RF classifiers generalize to new variants and capture the functional impact of a clinically challenging variant subset.

### Trained predictors uncover potential mechanisms of dysfunction for variants classified as “ambiguous” by AlphaMissense

AM classifies variants into three categories based on the calibrated pathogenicity score: likely pathogenic, likely benign, and ambiguous [21]. Variants falling within the ambiguous range represent cases where AM has low prediction confidence. We incidentally had function and trafficking measurements for 12 KCNQ1 variants classified as ambiguous (amb) by AlphaMissense (AM-amb) and used this set to test whether our RF classifiers could provide insight into the mechanisms underlying pathogenic ambiguity. Half of the variants exhibited I_peak_ LOF, with most trafficking normally, while effects on gating and kinetic parameters (V_1/2_, τ_act_, and τ_deact_) varied (**Fig. 4A**).

**Figure 4.**
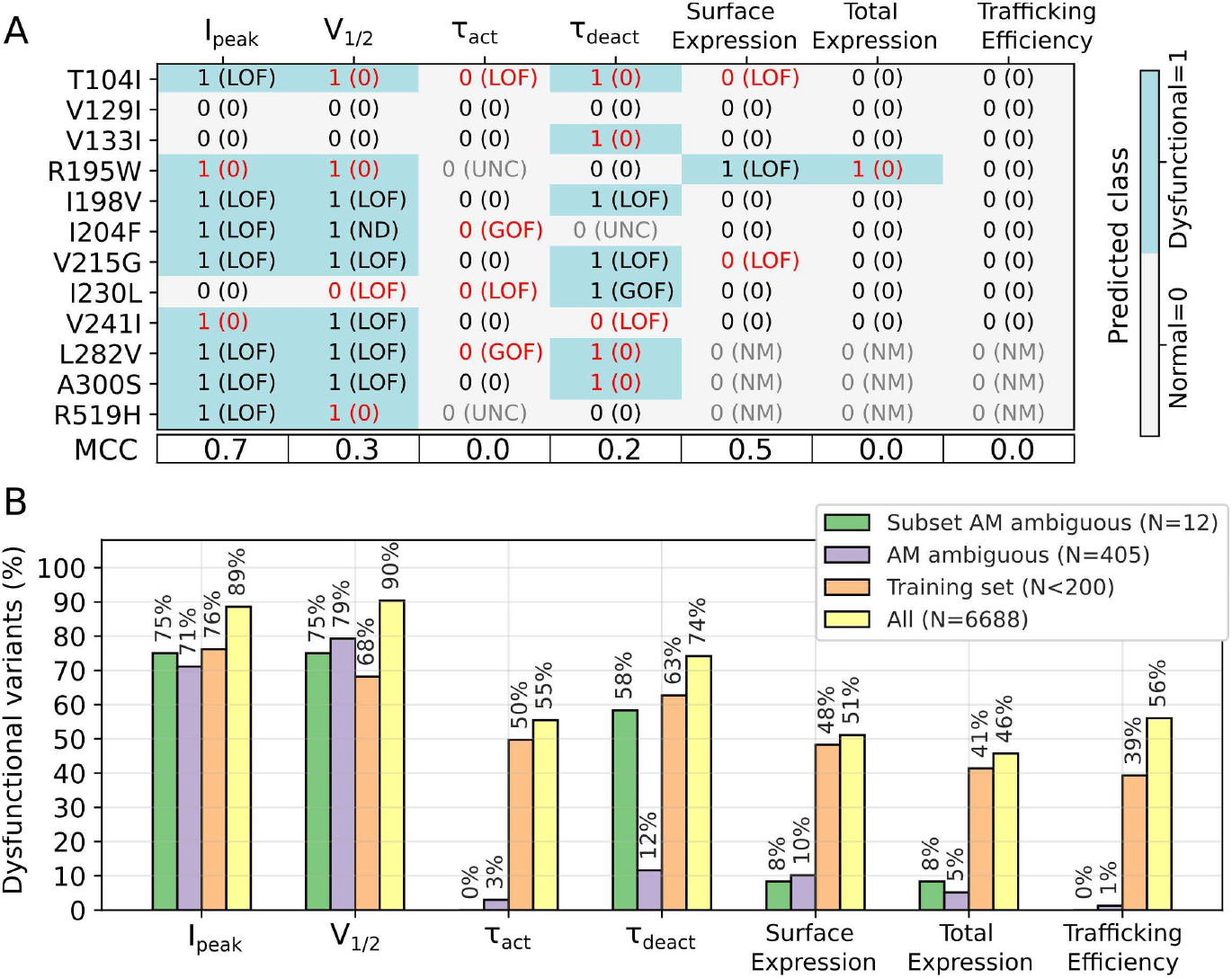
Predictions of AlphaMissense ambiguous (AM-amb) variants. (**A**) Heatmap where dysfunctional predictions are marked by blue cells. Text annotations are the same as in **Figure 3A**. Red text indicates the prediction does not match the true label and gray text indicates that the corresponding metric was either not measured (“NM”) or true label was uncertain (“UNC”). The model used for these predictions did not include these variants in the training set. MCC values are reported below respective predictions. UNC and NM measurements were not included in the MCC calculation. (**B**) Percentage of variants predicted to be dysfunctional. Bar colors represent different variant subsets, indicated by the figure legend. The total number of variants in each set is indicated in the legend (‘N’). RF classifiers provide insight for function and trafficking mechanisms of AM-amb variants.

Excluding the 12 AM-amb variants, we trained new RF classifiers as done for the ClinVar VUS in the previous section. Predictions and MCC values are reported in **Figure 4A**. Predictions mostly agreed with labeled classes (**Fig. 4A**, cells with *black* text). I_peak_ reflects the best model performance (MCC=0.7) on this test set, while τ_act_ and τ_deact_ performed the worst. All variants were predicted normal for τ_act_, however electrophysiological measurements indicated four were dysfunctional for τ_act_. Most variants were predicted to traffic normally, which agrees with the true labels.

Because trafficking efficiency was normal for the tested AM-amb variants (**Fig. 4A**), we sought to determine if this trend extends to all KCNQ1 AM-amb variants. Towards this, we used the trained RF classifiers to generate predictions for all KCNQ1 residue mutations spanning the protein range covered by our models and determined the ratio of variants predicted as dysfunctional for each measurement (**Fig. 4B**, *yellow*). While there was a disproportionately high percentage of variants predicted as dysfunctional for I_peak_ and V_1/2_, trafficking predictions were more evenly split between normal and dysfunctional variants (**Fig. 4B**, *yellow*). Therefore, it is possible that the RF classifiers are biased to predict dysfunctional I_peak_ and V_1/2_, but because of the 50-50 ratio of dysfunctional and normal variants in the trafficking training and “All” sets (**Fig. 4B**, *orange* and *yellow*), our predictions indicate that KCNQ1 AM-amb variants most likely traffic normally (**Fig. 4B**, *purple*). AM-amb predictions of time constants likewise indicate WT-like behavior (**Fig. 4B**, *purple*). However, the notably poor classification performance on the known AM-amb subset (**Fig. 4A**) makes us less inclined to draw conclusions on the general trend of τ_act_ and τ_deact_ for AM-amb variants. Further testing of more KCNQ1 AM-amb variants is needed to confirm these trends.

### Predictions clarify the interpretation of variants with uncertain experimental outcomes

To assign classification labels for variants from electrophysiological measurements, we would either use a cutoff value [24] or a statistical significance threshold [19]. In this study, we used both results to ensure the model learned biologically relevant trends on variants with statistically significant effects. For model training and validation, we excluded variants classified as “dysfunctional” by the cutoff threshold but with a non-significant p-value (p-value > 0.05). **Table 2** reports the count of these excluded “uncertain” variants determined by electrophysiology (**Figs. 1B-E**, *orange* spheres). The metric with the most uncertain variants was τ_act_ with 26 uncertain variants, whereas the other metrics had fewer than 10. The p-value threshold was not used for trafficking measurements because all variants outside the LOF and GOF cutoff thresholds show statistically significant differences from WT in the published set [19].

Due to the uncertainty in the true labels, this variant set was not used for performance evaluation. Rather, the computational predictions aided the interpretation of these variants. Towards this, we used the trained RF models to predict functional outcomes of the uncertain variants. To compare variant predictions and relative proximity to the cutoff and p-value thresholds, we plot electrophysiological measurements as in **Figures 1B-E** for the uncertain variants and color the point by predicted functional class (**Figs. 5A-D**). The electrophysiological measurements of variants predicted with normal WT-like function are near cutoff thresholds (**Figs. 5A-D**, *white* spheres near *dashed* lines). However, because a few of the variants predicted as dysfunctional were also near the cutoff threshold, adjusting the cutoff for classification could result in variant mislabeling. The majority of I_peak_ and V_1/2_ uncertain variants were predicted as dysfunctional, while half were predicted dysfunctional for time constants (**Figs. 5A-D**, *blue* spheres). If we had only used the measurement cutoff threshold to classify variants, these uncertain variants would have been labeled dysfunctional. In that case, our results would indicate higher prediction accuracy of uncertain I_peak_ and V_1/2_ variants than uncertain τ_act_ and τ_deact_. However, we also acknowledge here that there may be a prediction bias given the larger fraction of dysfunctional variants in the training data (**Fig. 4**, *orange*).

**Figure 5.**
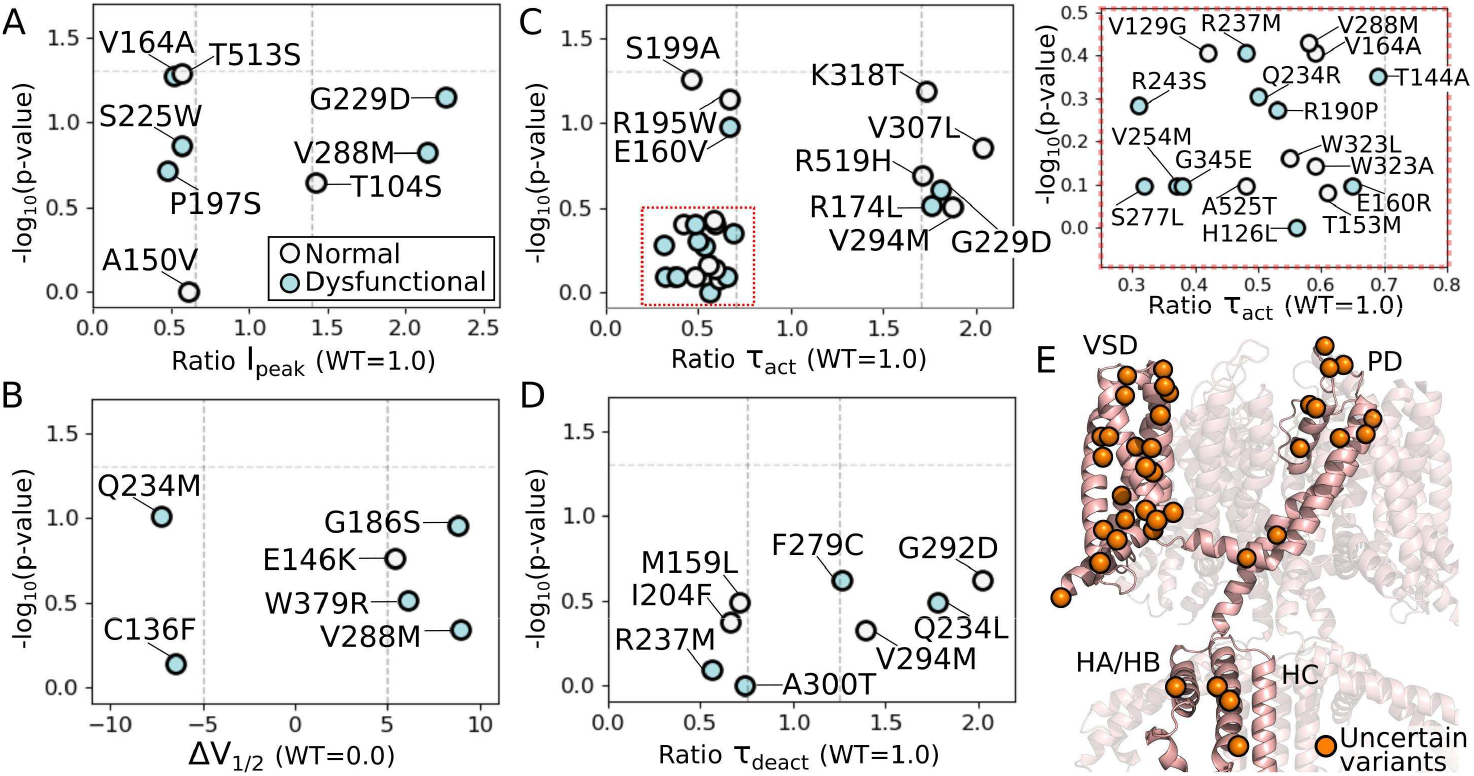
Mapping functional measurements and predictions for variants with experimentally-uncertain outcomes. (**A-D**) Functional measurements from electrophysiological recordings of the variants with uncertain class labels (**Figs. 1B-E**, *orange*). Colors correspond to RF classifier predictions, where white=normal WT-like function and blue=dysfunctional. Dashed lines mark the cutoff thresholds detailed in **Table 1**. A zoomed in view of the variants enclosed by the red dotted box in panel C are shown to the right of the panel. (**E**) Uncertain variants mapped onto a structural model of KCNQ1. VSD=voltage-sensing domain. PD=pore domain. HA, HB, HC=intracellular helical domains. Predictions from RF classifiers inform on the functional impact of variants with uncertain effects.

The uncertain variants are distributed throughout all domains of the KCNQ1 protein (**Fig. 5E**). For all uncertain variants, we systematically compared function and trafficking predictions with AM pathogenicity (**Fig. S3**). We analyzed the data from all predictions and, if available, true labels (**Fig. S3A**, cells with *black* and *UNC*). Variants that AM classified as benign (**Fig. S3B**, *benign* column) were mostly functional and trafficking normal (**Fig. S3A**, *white* cells), except for three variants with measured GOF I_peak_ (T153M, S199A, V294M) and two with predicted dysfunctional I_peak_ (V288M, G292D). V288M and G292D were both I_peak_ GOF based on electrophysiological measurements (**Fig. 5A**). This suggests that some AM benign variants might be GOF, a phenotype related to short QT syndrome [32-35]. While our model does predict all these GOF mutations as dysfunctional, future iterations of the model would benefit from including the gain- and loss-of-function distinctions. AM-amb variants (**Fig. S3B**, *ambiguous* column) are either I_peak_ LOF (I204F, R519H) or one of the trafficking measurements is not WT-like (R195W), but all other metrics are normal. Variants that AM classified as pathogenic (**Fig. S3B**, *pathogenic* column) had at least two metrics predicted as dysfunctional (**Fig. S3A**, *blue* cells). Different metrics were predicted dysfunctional for these variants, indicating they represent a functionally diverse set where each affects a distinct aspect of channel function, which our RF classifiers can differentiate.

### Global dysfunction and mistrafficking scores based on predictions of individual protein fitness metrics distinguish database-annotated variants

While individual predictions of the seven protein fitness metrics are widely informative, in order to capture broader patterns and intuitively interpret variant behavior, we determined “global scores” of dysfunction and mistrafficking using a weighted sum of the binary predictions. The predictions of the four electrophysiological metrics were summed to get the global score of dysfunction, where the weights were based on the proportion of determined measurements per metric (**Table 2**, *Normal* plus *LOF* plus *GOF*; excludes *Not determined* and *Uncertain*). These weights were 0.36, 0.2, 0.23, and 0.21 for I_peak_, V_1/2_, τ_act_, and τ_deact_, respectively. The relative weights assigned to each metric correspond to their expected importance for variant pathogenicity. I_peak_ is prioritized with the highest weight as its consistent measurement in electrophysiological experiments makes it the standard functional metric for clinical variant interpretation [36]. The predictions of the three trafficking metrics were summed to get the global score of mistrafficking, where weight was treated equally for each metric (0.33). We separated global scores for dysfunction and mistrafficking because in our previous work we found that many known pathogenic LOF variants trafficked normally but exhibited altered functional properties [19]. If we combined all protein fitness metrics into one score, the function and trafficking effects would cancel each other out, resulting in a non-informative metric.

To illustrate the utility of these scores, we compared the global dysfunction and mistrafficking scores to the clinical annotation for variants from ClinVar spanning the protein range covered by our model (residue positions 104-396 and 506-564). Because there were only four variants annotated as “benign” or “likely benign” within the covered range in ClinVar, we supplemented benign variants with missense variants reported in gnomAD. Variants with allele frequency greater than 1E-5 were considered too common to be causative for a rare disease like LQTS. This criterion identified 49 common variants, which we annotated as “likely benign.” The global scores per category indicate a clear and statistically significant difference between database-annotated benign (includes both “benign” and “likely benign”) and pathogenic (includes both “pathogenic” and “likely pathogenic”) variants (**Figs. 6A and 6B**). The scores do not distinguish between “benign” and “likely benign” (nor conversely between “pathogenic” and “likely pathogenic”), but the bar plots indicate that a variant with a global score of dysfunction below 0.4 is likely benign. We assigned numerical labels to each annotation (0 for “benign” or “likely benign” and 1 for “pathogenic” or “likely pathogenic”) and calculated the spearman rank correlation coefficient between annotation and global score of dysfunction (ρ=0.63) and annotation and global score of mistrafficking (ρ=0.48). The values indicate a moderate correlation between clinical annotation and global scores. We mapped the variants onto the KCNQ1 structure and colored the Cα atom of the residue by global scores (**Figs. 6C and 6D**). In the transmembrane region variants with “more benign” global score clustered at the VSD away from the PD, however pathogenic variants were distributed throughout the protein.

**Figure 6.**
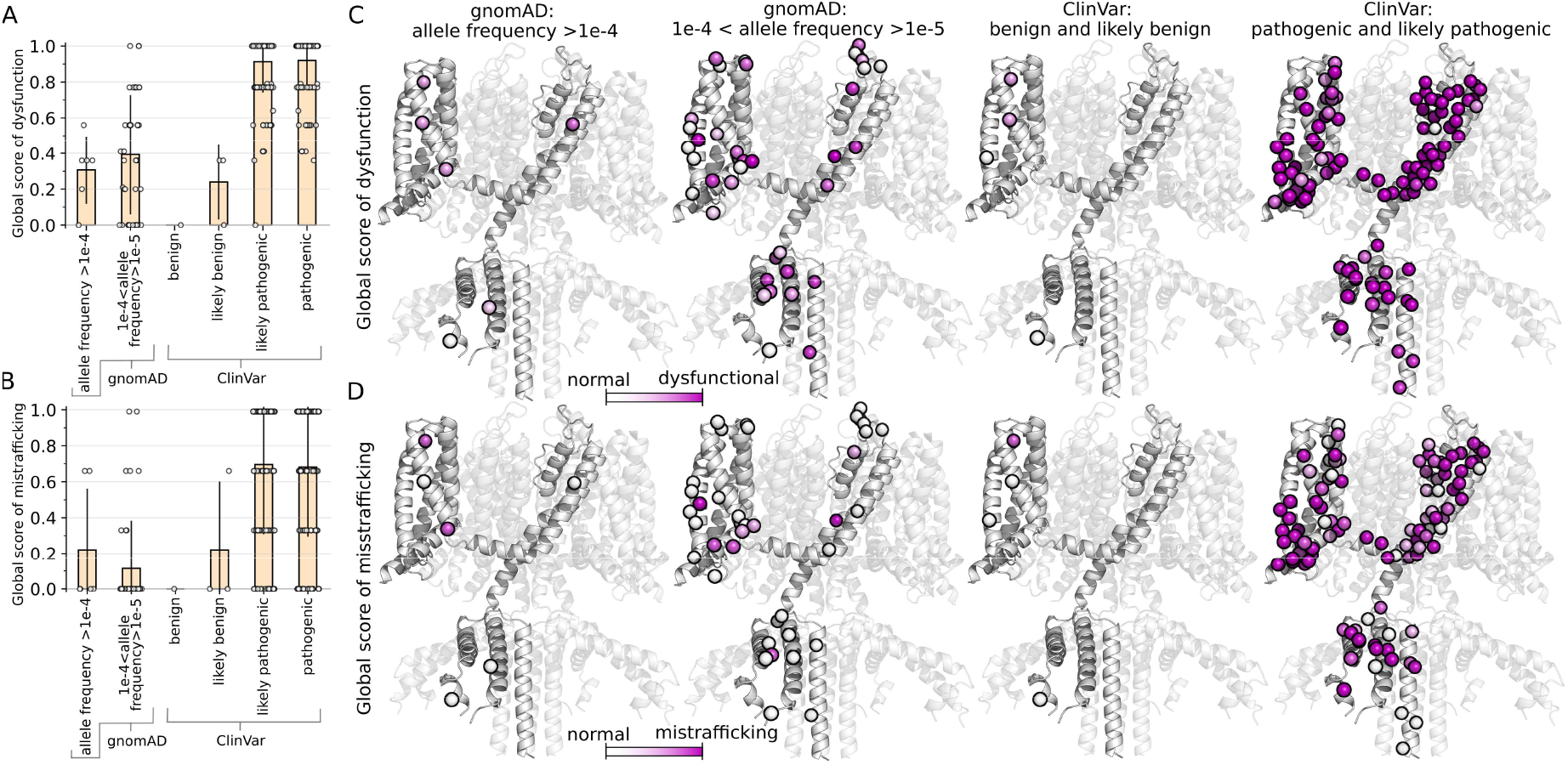
Global scores from protein fitness predictions to distinguish database-annotated benign and pathogenic variants. Bar plots depict mean and standard deviation of the global scores of (**A**) dysfunction and (**B**) mistrafficking of annotated variants from gnomAD and ClinVar. Scatter spheres represent variant scores which are calculated from the weighed sum of the function predictions and, separately, trafficking predictions. Values closer to 0 indicate the variant is WT-like. Variants from gnomAD with allele frequency greater than 1e-5 are considered as likely benign based on population frequency. However, these variants were also found in ClinVar and vary in assigned annotation, including benign/likely, uncertain, conflicting, and pathogenic/likely classifications. Some variants were reported in two or more sets in these bar plots if they have multiple entries in ClinVar or if they were found in both gnomAD and ClinVar databases. We kept duplicates to directly reflect database entries. (**B**,**C**) Structural mapping of gnomAD and ClinVar variants in panels A and B; Cα atoms are shown as spheres overlaid on KCNQ1. Domain notations are the same as in **Figure 1A**. Sphere colors correspond to the global score for the variant. Some residues had multiple variants in a set. The structure maps display the global score of one variant per residue; the score for the variant shown is automatically determined based on the order of variants in the list retrieved from ClinVar to avoid manual selection bias.

### Testing of all possible single-nucleotide missense variants spanning the S4 helix indicates accurate model performance

To evaluate model performance, we generated a focused set of all possible single-nucleotide missense variants at six codons within the S4 transmembrane helix, including positions with known disease-causing variants (S225, Q234, H240, G245) and positions without prior pathogenic reports (R228, R237) (the “S4” set; **Fig. 7A**). The positively-charged S4 helix is functionally important as it is the main voltage sensor of KCNQ1, where depolarization shifts the helix to the “up” conformation away from the intracellular side of the cell, a motion that is coupled to the opening of the pore domain [37]. Electrophysiological measurements are reported in **Figs. S4** along with RF model predictions (**Fig. S5A**). Some of the S4 variants were included in our training set, therefore we have duplicate electrophysiological measurements and additional trafficking measurements for those variants (**Fig. S5B**). We note here that our RF model would have seen these variants during training. Five mutations have identical amino acid variants resulting from different codon sequences (marked in **Fig. S4**). The RF models will predict the same label for these silent mutations. Consistent with the critical role of the S4 helix, AM predicts that none of the S4 variants are likely benign (**Fig. 7B**).

**Figure 7.**
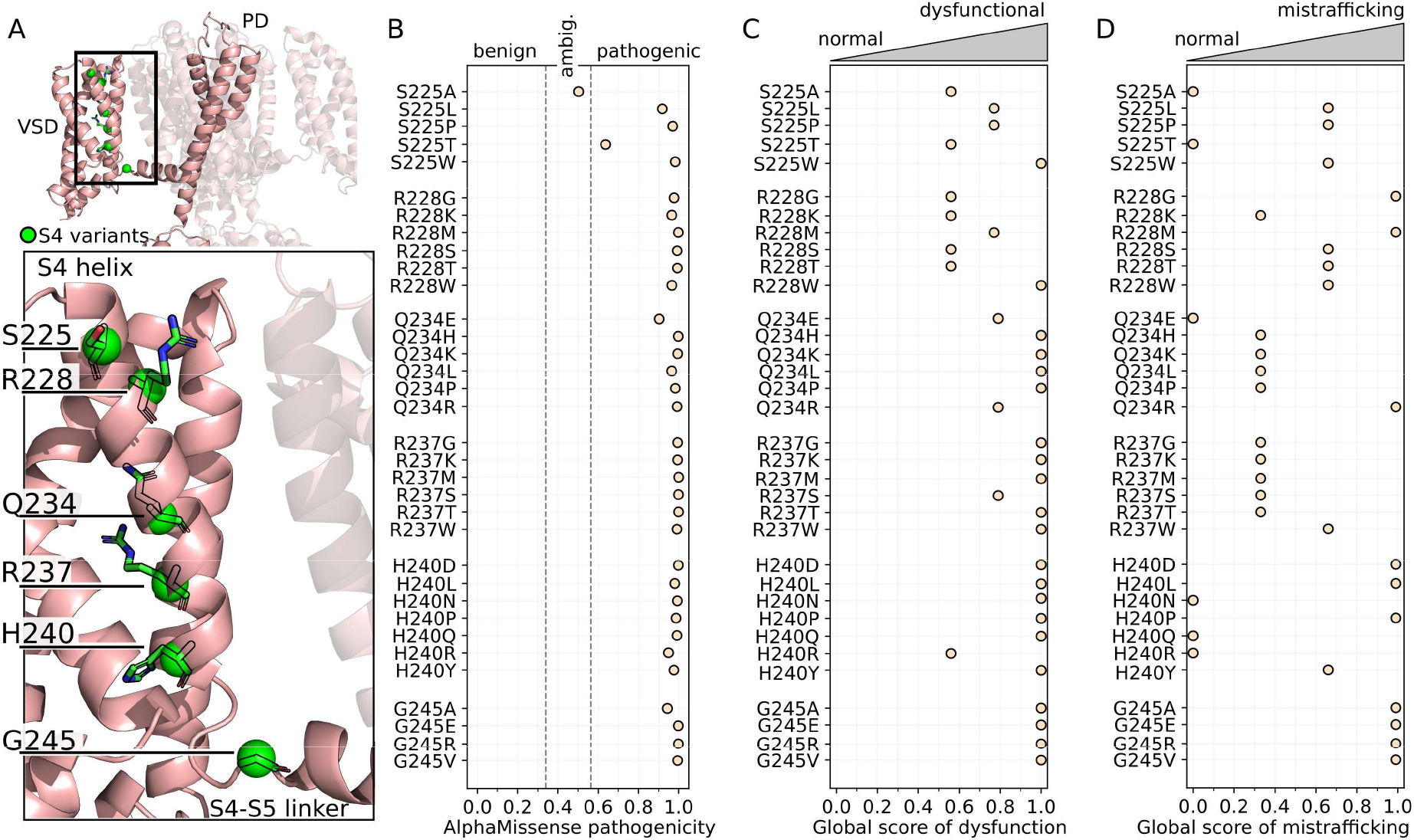
S4 variants structure mapping and predictions. (**A**) Structural mapping of S4 set variants. VSD=voltage-sensing domain. PD=pore domain. An enlarged view of the region enclosed by the black box displays the location of S4 variants in the VSD. Cα atoms are shown as spheres and side chains are shown with stick representation. (**B**) AM pathogenicity score of S4 variants. Class annotations from AM are indicated at the top of the plot. Global score of (**C**) dysfunction and (**D**) mistrafficking of S4 variants. Scores reflect the relative degree of dysfunction or mistrafficking for each variant, where a score closer to 0 indicates normal WT-like behavior. The developed global scores of dysfunction and mistrafficking distinguish between benign/likely and pathogenic/likely variants from the ClinVar and gnomAD databases.

The global scores of dysfunction (**Fig. 7C**) and mistrafficking (**Fig. 7D**) distinguished mechanistic trends for these residues. S225 did not tolerate bulky substitutions; dysfunction and mistrafficking scores were lowest for S225A and S225T, indicating that these substitutions maintained WT-like function (**Figs. 7C** and **7D**). In our previous experiments we found that S225L traffics at moderately higher levels than WT and was thermally stable [19]. In agreement with the higher dysfunction and mistrafficking scores, S225L and S225W are classified as likely pathogenic in ClinVar. R228 substitutions impacted trafficking but only partially disrupted function (**Figs. 7C** and **7D**). R228M and R228C were both supertraffickers, with all the trafficking measurements considerably above WT-like levels (**Table S1**). The side chain of R228 is oriented away from the VSD bundle in the VSD active and intermediate states (**Fig. 7A** and [38]), but faces the bundle interior in the VSD inactive state [37]. Therefore, substitutions are likely to affect VSD stability and the conformational transitions between functional states.

On the other hand, Q234 substitutions disrupted channel function, mostly without disrupting trafficking (**Figs. 7C** and **7D**). Previous experiments showed that Q234P, Q234M, and Q234C all traffic at WT-like levels [19, 20], and Q234M and Q234C both were thermally stable [19]. Q234P is classified as likely pathogenic in ClinVar, which our data indicates is due to altered function. Similarly, R237 substitutions consistently resulted in channel dysfunction, while trafficking was unaffected (**Figs. 7C** and **7D**). From our previous experiments, R237M and R237C traffic at moderately higher levels than WT and were thermally stable [19]. All R237 substitutions result in LOF I_peak_ (**Fig. S4A**), indicating this variant may be LQTS pathogenic, however there are no known disease-causing variants for the R237 codon.

H240 substitutions result in channel dysfunction, but half were predicted to traffic normally (**Figs. 7C** and **7D**). Mutation H240R was considered likely benign by our criteria from gnomAD (**Fig. 6**), which would agree with the lower global score of dysfunction for H240R compared to other substitutions (**Fig. 7C**) and prediction of WT-like trafficking (**Fig. 7D**). G245 does not tolerate any of the tested substitutions (**Figs. 7C, 7D**, and **S4**), indicating the importance of its location at the bend between the S4 and S4-S5 linker helices (**Fig. 7A**). Overall, these analyses demonstrate the utility of the global scores from our RF models in tandem with structural and clinical data to interpret variant effect on KCNQ1 function.

## Discussion

These results and our previous studies [19, 20, 24, 36] establish a comprehensive framework for interpreting KCNQ1 variants to enhance evaluation of rare or clinically ambiguous variants and support precision medicine efforts. In this study, we leveraged RF classifiers trained on biological data to predict the functional and trafficking effects of KCNQ1 missense variants. The value of the approach is illustrated across diverse use cases, including ClinVar VUS (**Fig. 3**), AM-amb variants (**Fig. 4**), and experimentally-uncertain variants (**Fig. 5**). Furthermore, we integrated individual predictions to distinguish database-annotated benign and pathogenic variants (**Fig. 6**) and systematically profiled variants within the functionally relevant S4 domain (**Fig. 7**). The independent classifiers allow for the distinction between variants that disrupt channel trafficking and those that alter channel function. The need for this approach is underscored by the heterogeneity in missense variant mechanisms associated with different disease states for membrane proteins like KCNQ1 [20, 39-41].

We attribute RF classifier improvements over the published Q1VarPredBio model [24] to the increased structural coverage of the protein, added features, and the change in model architecture. In testing different feature combinations, we found that using the 15 BioEvo+AM+MPNN+ESM features resulted in consistent performance for the seven predicted metrics (**Figs. 2** and **S1**). The individually trained classifiers reflect the inherent biological independence of each functional and trafficking measurement. Specifically, the four electrophysiology measurements are biologically independent of one another. Likewise, cell surface and total expression are contingent on post-translational processes or the transcriptional and translational machinery of the cell, respectively. A multi-class classifier may not capture the distinct patterns of each metric, reducing its ability to predict functional and trafficking outcomes separately. Individual models also allowed for targeted refinement of the training data by excluding variants with experimentally uncertain labels only from the relevant individual models. For example, we did not have to exclude all τ_act_ uncertain variants from the training set for the RF classifier that predicts I_peak_. This ensured that uncertainty in one measurement did not limit the ability of the model for another measurement to learn from that variant.

Published VEPs have different strengths and utilities, and therefore practitioners should evaluate models based on contextually-relevant variant subsets for their desired application when determining what predictive model to use [2, 42-44]. A recent assessment showed that although AM outperformed other methods overall, its performance varied widely across genes, highlighting the need for gene-specific variant classifiers [45]. Common concerns of VEPs include labeling errors in training data [2], inconsistent labels in different databases [46], and class imbalance leading to artificially inflated performance metrics [47, 48]. Variations in performance across models trained on different tasks suggest that no single VEP is universally appropriate for clinical classification, and evaluation criteria should match intended clinical or research use [2]. In this work, we report multiple cross-validation metrics across the full and focused variant subsets to provide a comprehensive evaluation of the RF classifiers. MCC is a stringent metric to evaluate imbalanced data, but its value is dependent on the classification threshold. Conversely Brier score uses predicted probabilities directly but can be misleading for very rare or very common outcomes [49, 50]. AUROC and AUPRC evaluate classification across varying thresholds. MCC and Brier scores indicate improved classification using the feature combinations with BioEvo features (**Fig. 2**), but for half of the prediction tasks, AUROC and AUPRC are consistently higher using AM alone than for the models with BioEvo features (**Fig. S1B** and **S1C**). Variants potentially in the AM training set were not excluded, which could inflate performance for the AM model. Evaluating ClinVar VUS (**Fig. 3**) and AM-amb (**Fig. 4**) subsets allowed comparison on variants likely unseen by AM. On these subsets, RF classifiers clearly outperformed the moderate predictive ability of AM, emphasizing the need to evaluate model performance using multiple metrics and variant subsets to minimize selection bias when comparing and deciding on what models to use for different tasks.

As with many biological applications, we worked with a small (N∼200) and imbalanced training set. While we used stratified shuffling to balance class labels for cross-validation, we did observe that predictions of all possible KCNQ1 missense mutations in the structured regions reflected the label distribution ratios in our training set (**Fig. 4B**). For I_peak_ and V_1/2_, about 90% of all KCNQ1 variants were predicted as dysfunctional, a considerably higher percentage than that for other metrics (**Fig. 4B**, *yellow*), consistent with the skewed distribution in the training set (**Fig. 4B**, *orange*, and **Table 1**). Trafficking predictions were more evenly distributed between classes (**Fig. 4B**, *yellow*), also reflected in the training set class label balance (**Fig. 4B**, *orange* and **Table 2**). While the high proportion of variants predicted as dysfunctional could reflect bias in the training data, it also mirrors biological constraints. Mutations in structured protein regions are generally less tolerated as they often result in misfolding and protein instability [19, 20, 41, 51]. In fact, most KCNQ1 variants reported in ClinVar within the examined sequence (residues 104-396 and 506-564) are classified as pathogenic or likely pathogenic (**Fig. 6**). The systematic analysis of S4 variants shows that our RF classifiers distinguish between dysfunctional and mistrafficking effects arising from different amino acid substitutions at a single position (**Fig. 7**).

To overcome data imbalance, we recommend the use of a general pathogenicity predictor like AM to select a minimal set of variants with expected class label balance and wide coverage throughout the protein prior to training protein fitness predictors. How many variants are considered “minimal” should be based on the constraints of the experimental methods used to measure the desired metrics in the wet lab. Baseline models like AM can predict metrics for a minimal variant set, extending to all mutations. Variants with low confidence, such as AM-amb, should be prioritized for testing, creating a feedback loop that focuses resources on the most informative, mechanistically ambiguous variants. This rational variant selection, similar to the strategy we used (see *Methods*) ensures models are trained using class-balanced, challenging variants.

Structure-aware VEPs demonstrate improved predictive performance over models without structural features [1, 2, 52]. We find that BioEvo features alone achieve strong performance for function and trafficking predictions (**Fig. 2**). To further evaluate feature impact, we compared feature importance of the 15 BioEvo+AM+MPNN+ESM features (**Fig. S6**). AM pathogenicity score and PSSM were consistently ranked in the topmost important features, followed by amino acid likelihoods from ProteinMPNN or ESM. Among biophysical features, distance from channel pore axis and local changes to hydrophobicity or polarizability were the most relevant. Conversely, features related to the nature of the amino acid were low in importance. The features from AM, ProteinMPNN, and ESM integrate extensive evolutionary, structural, and contextual information indicative of protein fitness. These features encode residue-level constraints, co-evolutionary relationships and mutational tolerance patterns learned from large-scale variant subsets across the proteome [21, 29, 31]. In contrast, structural features capture local modifications and subtle biophysical interactions, helping resolve protein- and metric-specific nuances. While combining these features yields robust KCNQ1 protein fitness classifiers, future VEP development may only require use of the most informative structural features with those from large-scale predictive models.

Overall protein fitness depends on complex factors, including collective dynamics and interactions with auxiliary subunits [7, 53-55]. A limitation of our approach is that all structural features were derived from a single WT channel conformation. We explored generating features from structural models representing different states (VSD up/down, pore open/closed) and also combined these state-dependent features to inform the model on channel dynamics. However, this approach resulted in limited impact on predictive performance, likely because the structural features we used do not differ substantially between structural models. For example, the distance from the channel pore axis will vary depending on pore state. However, there is only ∼2 Å distance shift of PD residues between the pore open and closed states [38], therefore most variants have the same distance from the channel pore axis, regardless of structural model. Channel dynamics shape protein fitness, as transitions between discrete conformational states guide activity. Missense mutations can disrupt this energy landscape [1]. Future VEPs could integrate variant-specific features derived from conformational ensembles that capture features like domain motions. While a handful of studies have used molecular dynamics for variant effect predictors [56-58], training data are often too sparse for sophisticated generative models. To overcome this, future work should leverage new methods to generate conformational ensembles [59] combined with protein fitness data from existing databases (i.e., MaveDB [60], DMS scans, ProteinGym [61]) and advanced artificial intelligence techniques to enable the development of highly accurate, dynamics-aware VEPS.

Whether a variant is a good candidate for therapeutic intervention either with a functional modulator, trafficking corrector, or both as in the case of the common cystic fibrosis deletion mutant [62], requires a comprehensive understanding of how the variant impacts the respective protein. While our RF classifiers perform well in distinguishing normal from dysfunctional or mistrafficking variants, the models do not distinguish between different forms of dysfunction, namely loss- and gain-of-function. Resolving the specific nature of the variant defect is key for clinical utility. For instance, a variant causing a reduction in trafficking would be an appropriate candidate for a therapeutic intervention that increases protein trafficking. However, if the same variant also causes an increase in current density, administering a trafficking corrector could lead to LQTS. Future efforts to develop protein fitness models should focus on differentiating LOF and GOF mutation effects on function and trafficking to provide better guidance for precision therapeutic interventions.

## Outlook and Conclusions

This work demonstrates the use of random forest classifier models to predict protein fitness metrics of the membrane protein KCNQ1 from a small, curated training set covering the structured domains of KCNQ1. Computational VEPs score the general pathogenicity but fundamentally lack the mechanistic detail crucial for informed clinical decision making, especially for complex membrane proteins like the KCNQ1 ion channel. This work shortens that critical gap. By training our classifiers on robust, experimentally derived KCNQ1 fitness metrics and integrating protein-specific with general predictive features (**Fig. 2**), we achieve reliable classification across distinct mechanistic categories on a challenging prediction task. This approach moves VEPs beyond ambiguous binary scoring, delivering a mechanistic predictor that pinpoints how a missense mutation leads to clinical disease. By evaluating the RF classifiers on focused subsets of variants (**Figs. 3-7**), we demonstrate the potential for contextual bias and the high predictive utility of the classifiers on specific training tasks, strongly supporting the recommendation that VEP performance must be assessed for problem- and system-specific applications [2, 46, 48, 63].

In the context of personalized medicine, understanding the specific molecular mechanism of dysfunction guides the development of precision treatment strategies tailored to an individual. Towards this, predictions for all structured-region KCNQ1 variants are provided as a publicly accessible resource. In addition, this study establishes a transferable paradigm where explicitly modeling biophysical and functional data is key to developing highly generalizable VEPs for membrane proteins, improving prediction and diagnosis across the entire ion channel superfamily. Overall, this work represents an advance in computational prediction tools of protein fitness, providing a framework that supports precision medicine through enhanced interpretation of variant effects across distinct mechanistic outcomes.

## Acknowledgments

This work was supported by NIH grant HL122010 (CRS, ALG, JM). ACCG is supported by T32 DK007061. E.W.B. was supported by the Integrated Training in Engineering and Diabetes T32 DK101003 and the National Institutes of Health F32 GM154455. JM acknowledges funding by the Deutsche Forschungsgemeinschaft (DFG, German Research Foundation) through SFB1423, project number 421152132. JM is supported by a Humboldt Professorship of the Alexander von Humboldt Foundation. The authors thank Alex Crowell for help with plasmid preparation and Katherine Clowes Moster for helpful discussions regarding trafficking data.

## Author Contributions

A.C.C.G. was the primary author of the manuscript, including developing the code for training, testing, and analyzing the classifier models, running benchmarks, generating figures, and writing the text. E.W.B. contributed code implementing the neural network for early benchmarks and provided guidance on model development, experimental design, data analysis, overall research direction, and contributed to manuscript draft revision. C.G.V. and E.D. performed variant functional testing, including measuring, analyzing, and collecting measurements. R.R.D. and J.M.D. conducted mutagenesis and heterologous expression studies. K.R.B. performed variant trafficking testing, including measuring, analyzing, and collecting measurements. T.S. manages the public code and data repository. K.V.L. provided intellectual input and guidance and contributed to manuscript draft revision. J.M., A.L.G., and C.R.S. oversaw the research and provided intellectual guidance. All authors assisted in and approve the revision and preparation of this manuscript for publication.

