## Supplemental Information for "Classification models distinguish functional and trafficking effects of KCNQ1 variants to enhance variant interpretation"

*[1]Dept. Chemistry, [2]Center for Structural Biology, [3]Dept. Biochemistry, [4]Institute of Chemical Biology, Vanderbilt University, Nashville, Tennessee, USA, [5]Dept. Medicine, Vanderbilt University Medical Center, Nashville, Tennessee, USA, [6]Dept. Pharmacology, Northwestern University Feinberg School of Medicine, Chicago, Illinois, USA, [7]Institute for Drug Discovery, Institute for Computer Science, Wilhelm Ostwald Institute for Physical and Theoretical Chemistry, University Leipzig, Leipzig, Germany, [8]Center for Scalable Data Analytics and Artificial Intelligence ScaDS.AI and School of Embedded Composite Artificial Intelligence SECAI, Dresden/Leipzig, Germany, [9]Center for Applied Artificial Intelligence in Protein Dynamics, Vanderbilt University, Nashville, Tennessee, USA*

**Content:**

Materials and Methods
Supplemental Figures S1-S6
Supplemental Tables S1-S2
Supplemental References

### Materials and Methods

**Mammalian cell culture.** CHO-K1 cells constitutively expressing human KCNE1 (designated CHO-KCNE1 cells) were generated as previously described [1] using the FLP-in™ system (Thermo Fisher Scientific, Waltham, MA, USA). Cells were grown in F-12 nutrient medium (GIBCO/Invitrogen, San Diego, CA, USA) supplemented with 10% fetal bovine serum (ATLANTA Biologicals, Norcross, GA, USA), penicillin (50 units/mL), streptomycin (50 µg/mL) at 37°C in 5% $CO_2$ and maintained under selection with hygromycin B (600 µg/mL).

**Plasmids and site-directed mutagenesis.** Full length cDNAs encoding wild type (WT) or variant human KCNQ1 (GenBank accession number AF000571) were engineered in the mammalian expression vector pIRES2-EGFP (BD Biosciences-Clontech, Mountain View, CA, USA) as previously described [1, 2]. This vector enabled expression of untagged channel subunits with fluorescent proteins as a means for tracking successful cell transfection. KCNQ1 variants were introduced into the WT coding sequence using Q5 Hot Start High Fidelity DNA polymerase (New England Biolabs, Ipswich, MA, USA). Mutagenic primers for each variant were designed using custom software (Q5 Designer; source code available at the Northwestern University Prism depository. Link: **https://prism.northwestern.edu/records/tcsgw-z4295** ). Variant KCNQ1 plasmid clones were screened as previously described [1]. For each variant plasmid, the complete KCNQ1 coding region, the IRES element and fluorescent protein cDNA were sequenced in their entirety by nanopore sequencing (Plasmidsaurus, South San Francisco, CA) and analyzed using a custom multiple sequence alignment tool (Multiple Sequence Iterative Comparator [MuSIC], available at https://doi.org/10.18131/h6hc6-n0j20). Endotoxin-free plasmid DNA was purified for clones with correct sequences (Nucleobond Xtra Maxi EF, Macherey-Nagel Inc.) and re-suspended in endotoxin-free water.

**Electroporation.** Plasmids encoding WT or mutant KCNQ1 channels were transiently transfected by electroporation using the Maxcyte STX system (MaxCyte Inc., Gaithersburg, MD, USA) [3]. CHO-KCNE1 cells grown to 70-80% confluence were harvested using 0.25% trypsin. A 500 µl aliquot of cell suspension was then used to determine cell number and viability using an automated cell_counter (ViCell, Beckman Coulter). Remaining cells were collected by gentle centrifugation (160 $\times$ g, 4 minutes), washed with 5 ml electroporation buffer (EBR100, MaxCyte Inc.), and re-suspended in electroporation buffer at a density of 100 x $10^6$ viable cells/ml. Each electroporation was performed using 100 µl of cell suspension. CHO-KCNE1 cells were electroporated with 10 µg of WT or mutant KCNQ1 cDNA. The DNA-cell suspension mix was transferred to an OC-100 processing assembly (MaxCyte Inc.) and electroporated using the preset CHO-PE (CHO_Protein Expression) protocol. Immediately after electroporation, 10 µl of recombinant human DNase (dornase alpha, Genentech, Inc., South San Francisco, CA) was added to the DNA-cell suspension and the entire mixture was transferred to a 35 mm tissue culture dish for a 30 min incubation at 37°C in 5% $CO_2$. Following incubation, cells were gently re-suspended in culture media, transferred to a T75 tissue culture flask and grown for 24 hours at 37°C in 5% $CO_2$. Following incubation, cells were harvested, counted, transfection efficiency determined by flow cytometry (see below), and then frozen in 1 ml aliquots at $1.8\times10^6$ viable cells/ml in liquid $N_2$ until used in experiments.

**Flow Cytometry.** Transfection efficiency was evaluated before freezing and prior to electrophysiological testing using a benchtop flow cytometer (CytoFLEX, Beckman Coulter, Brea CA, USA). Forward scatter (FSC), side scatter (SSC), green fluorescence (FITC) were recorded. FSC and SSC were used to gate single viable cells and to eliminate doublets, dead cells and debris. Ten thousand events were recorded for each sample. Nonelectroporated CHO-KCNE1 cells were assayed as a control for all parameters and used to set the gates for each experiment. A 488 nm laser was used to excite FITC, and the percentage of fluorescent cells was determined from the gated population.

**Electrophysiology.** The day before automated patch clamp recordings, electroporated cells were thawed, plated and incubated for 10 hours at 37°C in 5% $CO_2$. The cells were then grown overnight at 28°C in 5% $CO_2$ to increase channel expression at the plasma membrane. For the S4 variants, only an overnight growth incubation at 37°C was done. Prior to experiments, cells were passaged using 5% trypsin in cell culture media. Cell aliquots (500 µl) were used to determine cell number and viability by automated cell counting, and transfection efficiency by flow cytometry. Cells were then diluted to 200,000 cells/ml with external bath solution (see below) and allowed to recover for 60 minutes at 15°C while shaking on a rotating platform at 200 rpm.

Automated patch clamp experiments were performed using the Syncropatch 768 PE platform (Nanion Technologies, Munich, Germany) equipped with single-hole, 384-well recording S-Type chips (2-3 MΩ). For S4 variants, automated patch clamp experiments were performed using the Syncropatch 384i platform. Pulse generation and data collection were carried out with PatchController384 V.1.3.0 and DataController384 V1.2.1 software (Nanion Technologies, Munich, Germany). Whole-cell currents were recorded at room temperature in the whole-cell configuration, filtered at 3 kHz and acquired at 10 kHz. The access resistance and apparent membrane capacitance were estimated using built-in protocols. Access resistance was compensated by 80%. The external bath solution contained: 140 mM NaCl, 4 mM KCl, 2 mM $CaCl_2$, 1 mM $MgCl_2$, 10 mM HEPES, 5 mM glucose, pH 7.4. The internal solution contained: 60 mM KF, 50 mM KCl, 10 mM NaCl, 10 mM HEPES, 10 mM EGTA, 2 mM ATP-$K_2$, pH 7.2. Whole-cell currents were elicited from a holding potential of -80 mV using 2000 ms depolarizing pulses (from -80 mV to +60 mV in +10mV steps, every 10 secs) followed by a 2000 ms step to -30 mV to analyze tail currents and channel deactivation rate. Non-specific currents were eliminated by recording whole-cell currents before and after addition of the $I_{Ks}$ blocker JNJ-303 (2 µM). Whole-cell currents were not leak-subtracted. Only JNJ303-sensitive currents and recordings with seal resistance ≥ 0.5 GΩ, series resistance ≤ 20 MΩ, cell capacitance ≥ 1 pF for the S4 variants only, and voltage-clamp stability (defined as the standard error for the baseline current measured at the holding potential for all test pulses being <10% of the mean baseline current) were used in the final analysis.

**Electrophysiological data analysis.** Data were analyzed and plotted using DataController384 V1.8 (V3.2 for S4 variants) (Nanion Technologies, Munich, Germany), RStudio (Posit Software, Boston, MA, USA) for S4 variants, Excel (Microsoft Office 2013, Microsoft), SigmaPlot 2000 (Systat Software, Inc., San Jose, CA, USA) and Prism 8 (GraphPad Software, San Diego, CA). Additional custom semi-automated data handling routines were used for rapid analysis of current density and voltage-dependence of activation. Peak currents ($I_{peak}$) were recorded 1990 ms after the start of the voltage pulse, while tail currents were measured 10 ms after changing the membrane potential to -30 mV. The voltage-dependence of activation was calculated by fitting the normalized G-V curves with a Boltzmann function (tail currents measured at -30 mV). The voltage-dependence of activation was determined only for cells with mean current density greater than the background current amplitude. Rate of channel activation was determined by fitting activating currents at +60 mV to a single exponential function. Rate of channel deactivation was determined by fitting tail currents pre-activated with the +60 mV pulse to a single exponential function. Typical experiments compared five KCNQ1 variants to the WT channels assayed on the same plate with up to 64 replicate recordings. Properties of each KCNQ1 variant are presented relative to the WT channel assayed in parallel as percent peak current density measured at +60 mV, difference (Δ) in voltage-dependence of activation $V_{½}$, and the ratio of activation and deactivation time-constants (variant / WT). The number of cells used for each experimental condition is reported in the accompanying data.

**Trafficking measurements.** Trafficking measurements for variants used in this study were published in [4-6]. **Table S1** reports data for unpublished variants used in this study, which were measured with the same protocol as [4].

**S4 variants.** All possible single-nucleotide missense variants were designed at six codons within the S4 helix and a library (total of 48 variants) was generated of double stranded DNA cassettes encompassing codons 225 to 245 (Twist Biosciences, San Francisco, CA). From this library, pools of variants corresponding to each of the six selected codons were subcloned into a KCNQ1 expression plasmid using Gibson assembly. Individual plasmid clones were identified by Sanger sequencing and used in functional studies using automated voltage-clamp recording as detailed. Synonymous variants were included in the functional analyses, and nonsense variants were not. **Figure S4** reports electrophysiological measurements.

**Variant class assignment**. Both gain-of-function (GOF) and loss-of-function (LOF) variants were classified as dysfunctional. Some variants yielded too few cells or too low current for detectable measurement readouts (**Table 2,** *Not determined* column). These variants were classified as dysfunctional for model training. For training and validation, variants that fell outside the established WT-like threshold cutoff values but did not reach statistical significance (p-value > 0.05) were excluded (**Table 2,** *Uncertain* column). These were considered "uncertain" variants because their apparent dysfunction could not be confidently distinguished from measurement variability.

**Rational selection of variants for functional testing.** Variants of uncertain significance (VUS) were accessed from ClinVar and the online server of Q1VarPredBio [7] was used to generate function predictions. VUS set: From the list of VUS, 23 variants were manually selected. These variants were within the structured region of KCNQ1 covered by the model (see **Feature Generation**), had high Q1VarPredBio prediction confidence, evenly spanned the voltage-sensing domain (VSD) and pore domain (PD), and represented an approximately equal number of normal and dysfunctional predictions to create a balanced test set. The selected variants were functionally characterized as detailed in the experimental protocols. PD set: Noting the training set for Q1VarPredBio mostly consistent of variants from the VSD, a new set of variants spanning the S4-S5 linker through S6 was considered for electrophysiological recordings to include more variants from the PD in the training set. The number of variants for functional measurement was limited to a maximum of 24. In selecting the 24 variants, the strategy was to prioritize variants with predicted dysfunction probabilities near the middle range, meaning that they were neither clearly normal functioning (close to 0.0) nor clearly dysfunctional (close to 1.0), and located in regions of the protein with limited variant coverage. Twelve of the 24 variants were selected because at least one metric was predicted with a dysfunctional probability between 0.3 to 0.7. The other twelve were selected by identifying the 12 residues that were furthest in sequence distance from the residues covered by the training set. Of the 12, seven had mutations annotated in ClinVar as VUS and five were identified by predicting the functional outcomes of all possible mutations at residues in sparsely covered regions and selecting the mutation with the lowest prediction confidence from Q1VarPredBio. These 24 variants were functionally evaluated using electrophysiology as detailed in the experimental protocols above.

**Preliminary model benchmarks.** The Q1VarPredBio neural network architecture [7] was implemented in Python using PyTorch libraries. The published features and labels provided with the paper were used to replicate the reported MCC values for the original 125 variants. Q1VarPredBio is a multilayer perceptron (MLP) with 12 input features and four binary outputs corresponding to the four functional measurements. The Python implementation served as a baseline model to compare subsequent development. An updated set of features and labels including the additional 24 PD variants was generated and Q1VarPredBio was retrained. Performance was measured on 20 replicates of 5-fold cross-validation. In testing different feature combinations, using the 15 BioEvo+AM+MPNN+ESM features with the Q1VarPredBio MLP architecture [7] offered minor improvements to performance, but these were inconsistent in cross-validation folds and therefore collective MCC did not improve. Subsequently, different machine learning architectures were evaluated, and the RF classifiers consistently outperformed the artificial neural network used for Q1VarPredBio in cross-validation studies using the full training set and on smaller subsets, leading us to select the RF architecture for this work.

**Feature generation.** PDB 8SIK [8] was modified for feature generation. 8SIK represents the voltage sensor up conformational state with the pore domain closed The cryoEM model was aligned to the a1q1.pdb model used for Q1VarPredBio and the ModLoop web server [9, 10] was used to model the missing loops at residues 219-222 for chain A of 8SIK. The unstructured N- and C-termini of KCNQ1 and the region from the HA/HB helices to the HC helix were missing from the cryoEM model. These regions are largely disordered, and therefore were excluded from the model. Calmodulin and the calcium ions included in the cryoEM model were removed. This left only the KCNQ1 tetramer covering the structured regions from residues 104-396 and 506-564, which includes KCNQ1 transmembrane and cytosolic domains (**Fig. 1**). Model performance using features derived from the other available cryoEM structures [8, 11] or combinations of multiple cryoEM structures was tested. However, using the features derived from 8SIK-only consistently performed best (data not shown).

A similar BCL (Biology and Chemistry Library) protocol as detailed in [7] was employed to generate the BioEvo features using the new KCNQ1 structural model ("input_8sik_clean.pdb"). These features were extensively described in [7]. In brief, they include: hydrophobicity and polarizability of the mutant amino acid, functional density of polarizability and hydrophobicity within the local environment of the mutated residue, distance from the channel pore axis, burial on the membrane, the change in number of hydrogen acceptor and donor sites between the WT and mutated amino acid, the change of volume, and the change in likelihood of amino acid substitution at the mutation site as determined by searching the NCBI non-redundant sequence database [12]. Functional density accounts for the impact of the mutation on the local environment and is calculated by averaging the property of interest for residues within a given radius of the site of the variant and weighing by the inverse of the residue distance from the site of mutation [13]. KCNQ1 relies on hydrophobic residues to anchor and stabilize its transmembrane domains within the lipid bilayer, a property that is critical for proper folding and function. Hydrophobicity was quantified using the published hydrophobicity scale [14]. Polarizability is a function of the residue index of refraction, molecular weight, density, and number of atoms to relate to molar refractivity [15]. The radii used to define local environment for hydrophobicity and polarizability are the same as in Q1VarPredBio [7]. The rationale for including these features is that amino acids in similar local environments tend to share physicochemical properties; thus, substitutions that differ from their surroundings are more likely to perturb normal protein function. These 12 features were generated using the following command-line prompt to execute the BCL:

```
bcl.exe descriptor:GenerateDataset -source "ProteinMutationsDirectory(./, key file=./keys.txt, aa class=AAComplete, mutation extension=.model_data.csv, suffix=.pdb, add self mutation fraction=0.0)" -output temp_bcl_features_kcnq1.csv -feature_labels features_combined.obj -result_labels resultslqts_combined.obj -id_labels 'Combine (MutationId)'
```

The current implementation of the BCL `descriptor:GenerateDataset` function prints placeholder “labels” for the variants in the output file with the generated features. These were manually deleted and not used.

The AlphaMissense pathogenicity scores were obtained from the published AlphaMissense predictions [16]. The ProteinMPNN [17] and ESM [18, 19] amino acid probabilities were generated using the ProteinMPNNProbabilitiesMetric and the PerResidueEsmProbabilitiesMetric, respectively, movers implemented in Rosetta [20]. The following prompt executes the movers, where `$ROSETTA` should point to the location of Rosetta program files:

```
$ROSETTA/main/source/bin/rosetta_scripts.pytorchtensorflow.linuxgccrelease -s $PDB -parser:protocol predict_probs.xml -out:file:score_only score.sc
```

The AlphaMissense, ProteinMPNN, and ESM features were appended to the BioEvo features using stand-alone Python scripts to process the data. The full set of 15 BioEvo+AM+MPNN+ESM features used for the RF classifiers is found in “features_8sik_clean.w_am_mpnn_esm.csv”.

**RF classifiers.** The random forest (RF) classifiers were implemented using the scikit-learn library in Python (“main.py”). In total we trained 21 RF classifiers: seven predicted measurements ($I_{peak}$, $V_{1/2}$, $\tau_{act}$, $\tau_{deact}$, cell surface expression, total expression, and trafficking efficiency) times three feature combinations (BioEvo, BioEvo+AM, and BioEvo+AM+MPNN+ESM). All procedures were executed using a YAML configuration file; example YAML files are in the sample_input_yaml folder. The script uses the function and trafficking measurements from experiments (“kcnq1_all_measurements.csv”) to generate class labels based on the cutoff thresholds (**Table 1** and p-value cutoff as detailed in the main text). The scikit-learn StandardScaler was applied to normalize input features. Grid search with cross-validation was used to find the optimal parameters for each RF classifier. Aware of the possibility of overfitting, tree depth was restricted while allowing a larger number of trees in the parameters grid. A custom scoring function was used to evaluate model performance during hyperparameter tuning. The custom scoring function combined the Matthews correlation coefficient (MCC) at the default 0.5 classification threshold with Brier score, with weights of 0.7 and 0.3, respectively. The weights were empirically determined to balance classification performance while accounting for model calibration to guide the search towards models with strong discriminative ability and acceptable probability metrics. As an example, **Table S2** lists the number of trees and forest depth of the RF classifiers trained using the BioEvo+AM+MPNN+ESM features. The optimized hyperparameters for each RF classifier are provided in “saved_models/out_rfc._hyperparams.joblib”.

RF classifier cross-validation was performed using the optimized hyperparameters with 20 replicates of 5-fold cross-validation and stratified sampling. A scikit-learn Pipeline with a StandardScaler followed by the RF classifier was implemented such that during model training, the scaler computes the mean and standard deviation from the training set to apply to the training and validation splits consistently. This ensures that predictions on the validation set are made on features normalized based on the training set. For each predicted measurement, a for loop performs model fitting and predictions for the three feature combinations using the same training/validation splits such that the validation metrics per fold are comparable across feature combinations. The optimal decision threshold for binary classification was selected by maximizing the MCC across a tested range of probability thresholds from 0.1 to 0.9 in increments of 0.01. Cross-validation variants, labels, predictions, and classification threshold were saved to “saved_models/out_rf.[measurement].joblib” and were used to generate **Figures 2** and **S1**.

**RF classifier evaluation.** Model performance was evaluated using MCC, Brier score, AUPRC, and AUROC. Matthews correlation coefficient (MCC) measures classification performance by considering all elements of the confusion matrix, making it a suitable metric for imbalanced data. MCC values range from -1.0 to 1.0, where 1.0 indicates perfect agreement between the two sets of binary data, 0.0 indicates no correlation, and -1.0 indicates a perfectly negative correlation. Brier score evaluates predictive performance by calculating the mean squared difference between the predicted probabilities and the true labels, providing a measure of prediction quality. Brier scores range from 0.0 to 1.0, where scores closer to 0.0 indicate alignment between predicted probabilities and labels, one metric used to assess model calibration. Area under the precision-recall curve (AUPRC) and area under the receiver operating characteristic curve (AUROC) were calculated from curves that evaluate the classifier predictions at various probability thresholds. They range from 0.0 to 1.0, where values closer to 1.0 indicate that the classifier more effectively identifies positive and negative labels across continuous thresholds.

**Saved models.** Models were trained as detailed in the text and the model states were saved.

- The model trained on all variants is saved_models/trained_on_all/save_model_state.rfc.biophys_evol_am_mpnnesm.*.joblib
- The model trained excluding AlphaMissense ambiguous variants is saved_models/test_ambig/save_model_state.rfc.biophys_evol_am_mpnnesm.excl_ambiguous.*.joblib
- All variant predictions are in kcnq1_predictions.hdf5
- The code to read the predictions file and calculate and plot global score weights as done for **Figure 6** is additional_analyses_scripts/x_analyze_global_score.py.
- In “list_variant_subsets/” we include the variant lists from ClinVar and gnomAD and the list of all KCNQ1 variants annotated as ambiguous by AlphaMissense

**Data and code repository**: https://github.com/meilerlab/kcnq1-function-trafficking.git

### Supplemental Figures

#### Figure S1

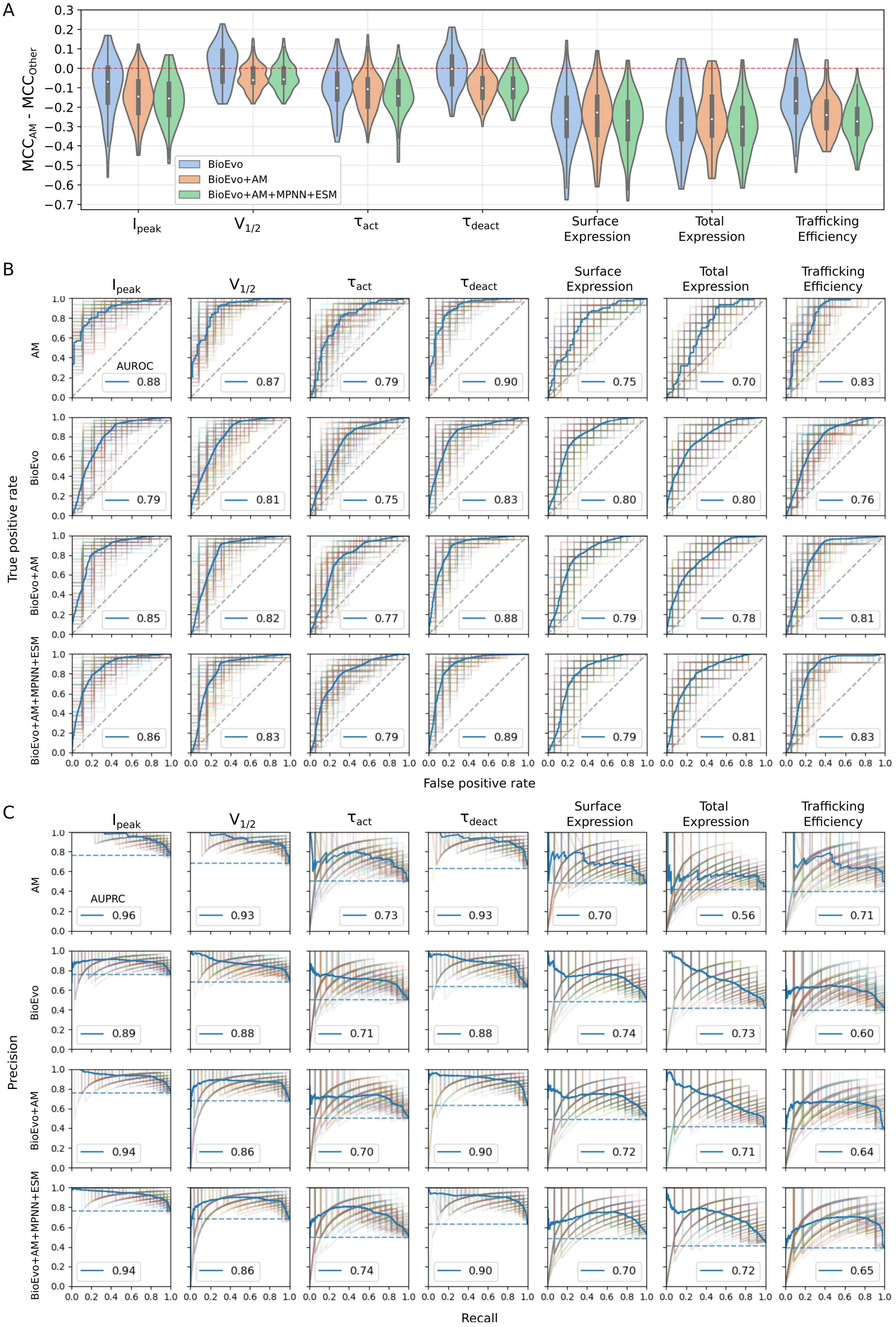


**Figure S1. Additional cross-validation metrics.** These metrics were collected from the same data shown in **Figure 2**. (**A**) Difference MCC per validation set. $MCC_{AM}$ uses the AM pathogenicity score directly as a predictive value of the metric (**Fig. 2A**, *blue*). $MCC_{Other}$ uses the predictions from our RF classifiers with respective feature set as indicated in the legend corresponding with **Figure 2A.** The same variants were included in the validation set of each fold for each model, so the per-fold validation metrics are comparable across models. (**B**) Receiver operating curves and (**C**) precision-recall curves for the same data as in **Figure 2**. There are 100 partially transparent curves representing each validation set. The solid blue curve in each plot includes all the validation sets combined. Area under the receiver operating curve (AUROC) and area under the precision recall curve (AUPRC) of the blue curve is reported in the panel legend.

**Figure S2**

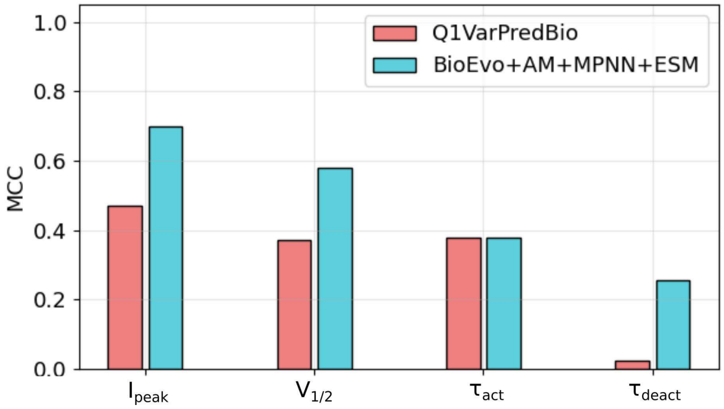


**Figure S2. Test set performance for two models.** Matthews correlation coefficient calculated from function predictions of a subset of ClinVar VUS (**Fig. 3**). Predictions were either from the published Q1VarPredBio [7] or the RF classifiers in this work. Except for $\tau_{act}$, RF classifiers improve function predictions on this subset of VUS.

**Figure S3**

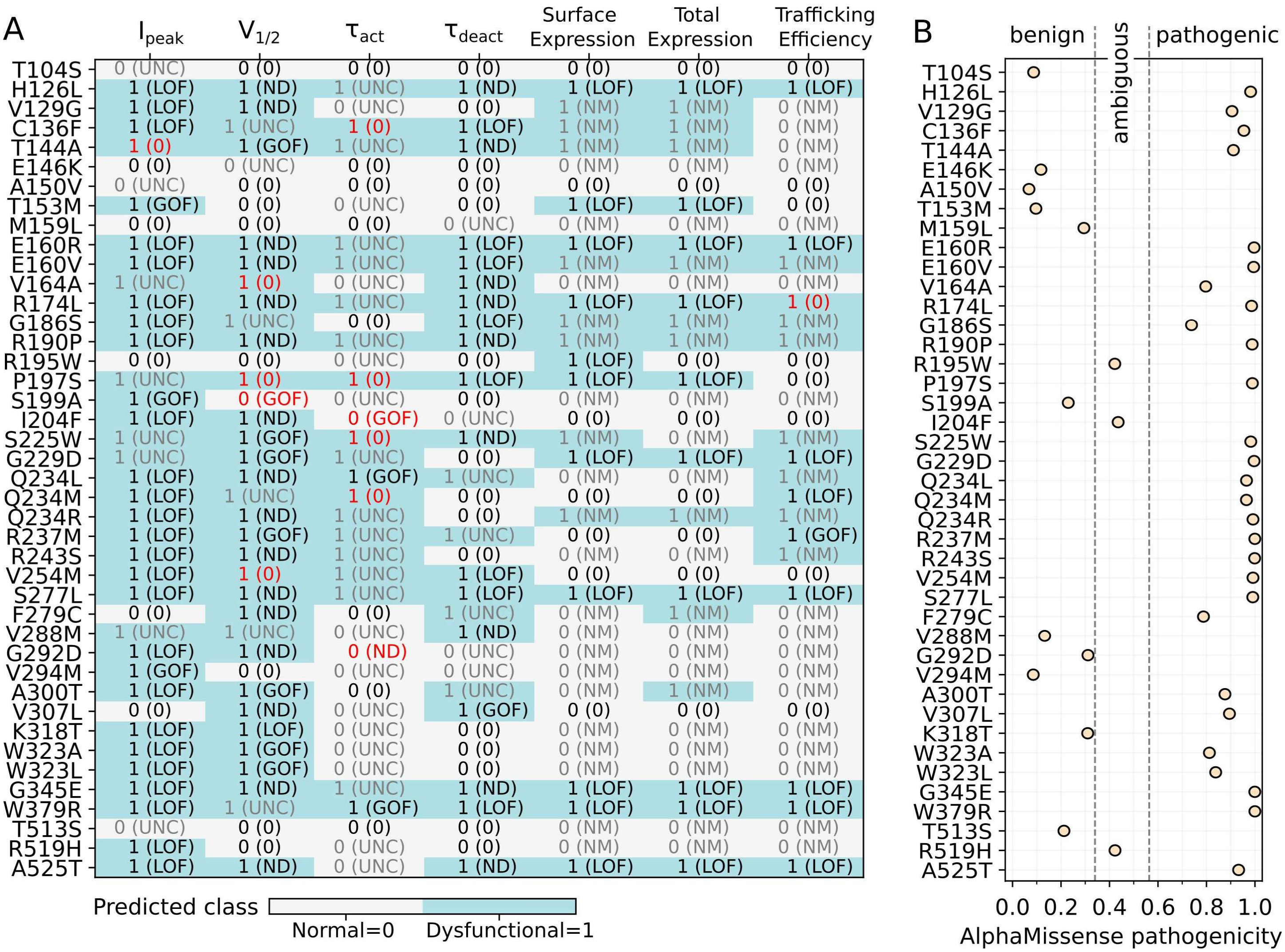


**Figure S3. Model predictions for experimentally-uncertain variants.** The variants in gray text marked "UNC" for $I_{peak}$ correspond to the variants labeled in **Figure 5A**, and similarly for the variants marked "UNC" for $V_{1/2}$ (**Fig. 5B**), $\tau_{act}$ (**Fig. 5C**), and $\tau_{deact}$ (**Fig. 5D**). (**A**) Heatmap where variants predicted as dysfunctional are marked by blue cells. Text annotations include the predicted class (normal=0, dysfunctional=1) and the true label in parentheses. LOF, GOF, and ND variants were considered "dysfunctional" for classification labels. Black text in the cells indicates the assigned class label and prediction agree. Red text indicates the prediction does not match the label. Gray text indicates that the experimental value (true label) for the variant was either uncertain ("UNC") or not measured ("NM"). (**B**) AM pathogenicity score for corresponding variants in panel A. Class annotations from AM are indicated at the top of the plot.

**Figure S4**

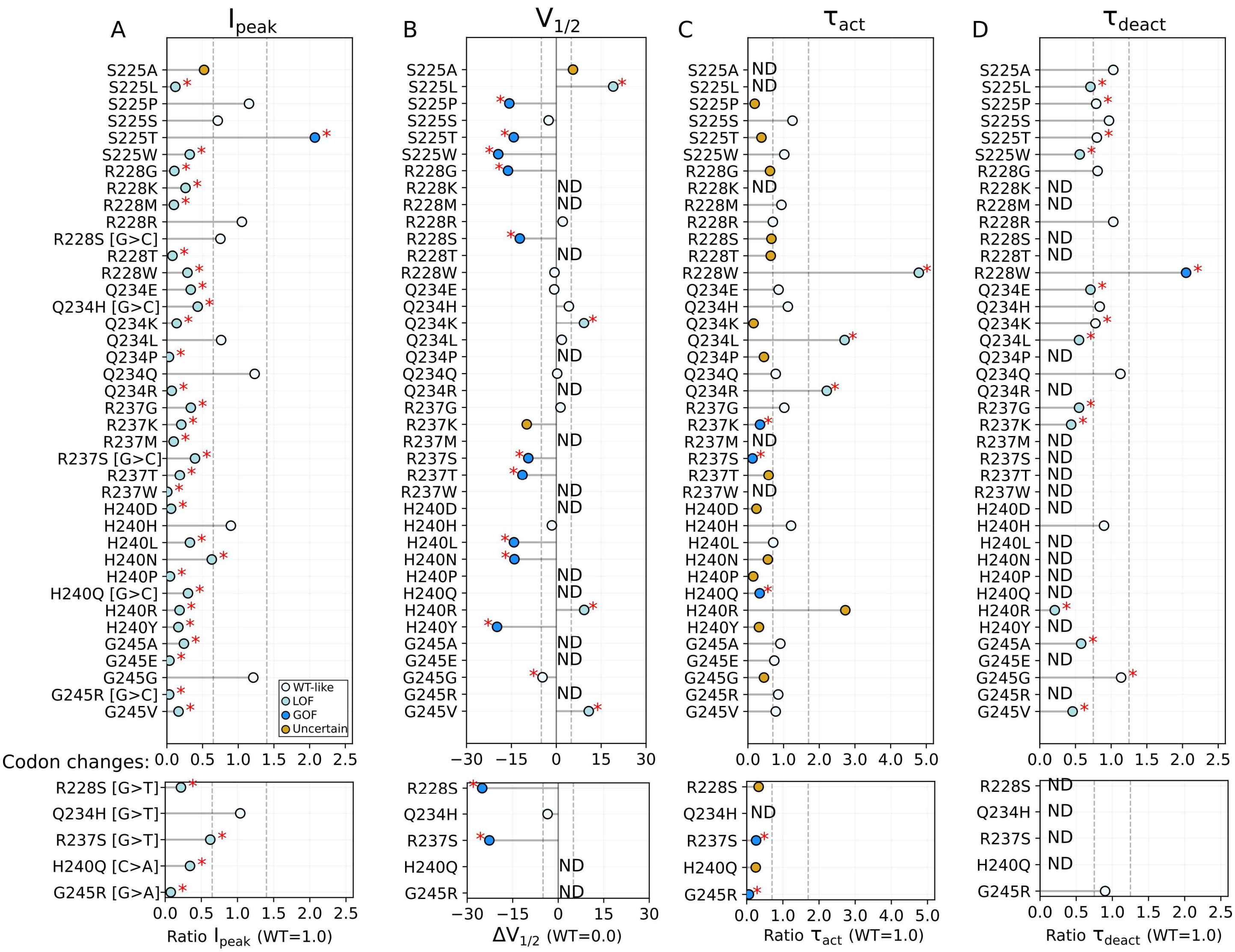

**Figure S4. Function measurements of S4 set variants.** (**A-D**) Variant measurements from electrophysiological recordings scaled to WT. Dashed lines denote cutoff thresholds for classification (**Table 1**). Circle marker colors correspond to assigned class labels as follows: cyan=LOF, white=WT-like normal, blue=GOF, and orange=uncertain. The red asterisk indicates the measured value was statistically significant (p-value<=0.05). Variants with “Not determined” measurements are marked “ND.” The variants with codon changes are marked in panel A and carry through to panels B-D. In the top panel, the reported measurement used the variant expressed using the G>C codon. In the bottom panel, the measurement used the variant expressed with the specified codon.

#### Figure S5

A

| | $I_{peak}$ | $V_{1/2}$ | $\tau_{act}$ | $\tau_{deact}$ | Surface Expression | Total Expression | Trafficking Efficiency |
|---|---|---|---|---|---|---|---|
| S225A | 1 (UNC) | 1 (UNC) | 0 (ND) | 0 (0) | 0 (NM) | 0 (NM) | 0 (NM) |
| S225L | 1 (LOF) | 1 (LOF) | 0 (ND) | 1 (LOF) | 1 (NM) | 0 (NM) | 1 (NM) |
| S225P | 1 (0) | 1 (GOF) | 0 (UNC) | 1 (0) | 1 (NM) | 0 (NM) | 1 (NM) |
| S225S | 0 (0) | 0 (0) | 0 (0) | 0 (0) | 0 (NM) | 0 (NM) | 0 (NM) |
| S225T | 1 (GOF) | 1 (GOF) | 0 (UNC) | 0 (0) | 0 (NM) | 0 (NM) | 0 (NM) |
| S225W | 1 (LOF) | 1 (GOF) | 1 (0) | 1 (LOF) | 1 (NM) | 0 (NM) | 1 (NM) |
| R228G | 1 (LOF) | 1 (GOF) | 0 (UNC) | 0 (0) | 1 (NM) | 1 (NM) | 1 (NM) |
| R228K | 1 (LOF) | 1 (ND) | 0 (ND) | 0 (ND) | 0 (NM) | 1 (NM) | 0 (NM) |
| R228M | 1 (LOF) | 1 (ND) | 0 (0) | 1 (ND) | 1 (NM) | 1 (NM) | 1 (NM) |
| R228R | 0 (0) | 0 (0) | 0 (0) | 0 (0) | 0 (NM) | 0 (NM) | 0 (NM) |
| R228S [G>C] | 1 (0) | 1 (GOF) | 0 (UNC) | 0 (ND) | 0 (NM) | 1 (NM) | 1 (NM) |
| R228T | 1 (LOF) | 1 (ND) | 0 (UNC) | 0 (ND) | 0 (NM) | 1 (NM) | 1 (NM) |
| R228W | 1 (LOF) | 1 (0) | 1 (LOF) | 1 (GOF) | 1 (NM) | 0 (NM) | 1 (NM) |
| Q234E | 1 (LOF) | 1 (0) | 1 (0) | 0 (LOF) | 0 (NM) | 0 (NM) | 0 (NM) |
| Q234H [G>C] | 1 (LOF) | 1 (0) | 1 (0) | 1 (0) | 0 (NM) | 0 (NM) | 1 (NM) |
| Q234K | 1 (LOF) | 1 (LOF) | 1 (UNC) | 1 (0) | 0 (NM) | 0 (NM) | 1 (NM) |
| Q234L | 1 (0) | 1 (0) | 1 (LOF) | 1 (LOF) | 0 (NM) | 0 (NM) | 1 (NM) |
| Q234P | 1 (LOF) | 1 (ND) | 1 (UNC) | 1 (ND) | 0 (NM) | 0 (NM) | 1 (NM) |
| Q234Q | 0 (0) | 0 (0) | 0 (0) | 0 (0) | 0 (NM) | 0 (NM) | 0 (NM) |
| Q234R | 1 (LOF) | 1 (ND) | 1 (LOF) | 0 (ND) | 1 (NM) | 1 (NM) | 1 (NM) |
| R237G | 1 (LOF) | 1 (0) | 1 (0) | 1 (LOF) | 0 (NM) | 0 (NM) | 1 (NM) |
| R237K | 1 (LOF) | 1 (UNC) | 1 (GOF) | 1 (LOF) | 0 (NM) | 0 (NM) | 1 (NM) |
| R237M | 1 (LOF) | 1 (ND) | 1 (ND) | 1 (ND) | 0 (NM) | 0 (NM) | 1 (NM) |
| R237S [G>C] | 1 (LOF) | 1 (GOF) | 1 (GOF) | 0 (ND) | 0 (NM) | 0 (NM) | 1 (NM) |
| R237T | 1 (LOF) | 1 (GOF) | 1 (UNC) | 1 (ND) | 0 (NM) | 0 (NM) | 1 (NM) |
| R237W | 1 (LOF) | 1 (ND) | 1 (ND) | 1 (ND) | 1 (NM) | 0 (NM) | 1 (NM) |
| H240D | 1 (LOF) | 1 (ND) | 1 (UNC) | 1 (ND) | 1 (NM) | 1 (NM) | 1 (NM) |
| H240H | 0 (0) | 0 (0) | 0 (0) | 0 (0) | 0 (NM) | 0 (NM) | 0 (NM) |
| H240L | 1 (LOF) | 1 (GOF) | 1 (0) | 1 (ND) | 1 (NM) | 1 (NM) | 1 (NM) |
| H240N | 1 (LOF) | 1 (GOF) | 1 (UNC) | 1 (ND) | 0 (NM) | 0 (NM) | 0 (NM) |
| H240P | 1 (LOF) | 1 (ND) | 1 (UNC) | 1 (ND) | 1 (NM) | 1 (NM) | 1 (NM) |
| H240Q [G>C] | 1 (LOF) | 1 (ND) | 1 (GOF) | 1 (ND) | 0 (NM) | 0 (NM) | 0 (NM) |
| H240R | 1 (LOF) | 1 (LOF) | 0 (UNC) | 0 (LOF) | 0 (NM) | 0 (NM) | 0 (NM) |
| H240Y | 1 (LOF) | 1 (GOF) | 1 (UNC) | 1 (ND) | 0 (NM) | 1 (NM) | 1 (NM) |
| G245A | 1 (LOF) | 1 (ND) | 1 (0) | 1 (LOF) | 1 (NM) | 1 (NM) | 1 (NM) |
| G245E | 1 (LOF) | 1 (ND) | 1 (0) | 1 (ND) | 1 (NM) | 1 (NM) | 1 (NM) |
| G245G | 0 (0) | 0 (0) | 0 (UNC) | 0 (0) | 0 (NM) | 0 (NM) | 0 (NM) |
| G245R [G>C] | 1 (LOF) | 1 (ND) | 1 (0) | 1 (ND) | 1 (NM) | 1 (NM) | 1 (NM) |
| G245V | 1 (LOF) | 1 (LOF) | 1 (0) | 1 (LOF) | 1 (NM) | 1 (NM) | 1 (NM) |

Codon changes:

| | $I_{peak}$ | $V_{1/2}$ | $\tau_{act}$ | $\tau_{deact}$ | Surface Expression | Total Expression | Trafficking Efficiency |
|---|---|---|---|---|---|---|---|
| R228S [G>T] | 1 (LOF) | 1 (GOF) | 0 (UNC) | 0 (ND) | 0 (NM) | 1 (NM) | 1 (NM) |
| Q234H [G>T] | 1 (0) | 1 (0) | 1 (ND) | 1 (ND) | 0 (NM) | 0 (NM) | 1 (NM) |
| R237S [G>T] | 1 (LOF) | 1 (GOF) | 1 (GOF) | 0 (ND) | 0 (NM) | 0 (NM) | 1 (NM) |
| H240Q [C>A] | 1 (LOF) | 1 (ND) | 1 (UNC) | 1 (ND) | 0 (NM) | 0 (NM) | 0 (NM) |
| G245R [G>A] | 1 (LOF) | 1 (ND) | 1 (GOF) | 1 (0) | 1 (NM) | 1 (NM) | 1 (NM) |

Predicted class: Normal=0, Dysfunctional=1

B

Replicate:

| | $I_{peak}$ | $V_{1/2}$ | $\tau_{act}$ | $\tau_{deact}$ | Surface Expression | Total Expression | Trafficking Efficiency |
|---|---|---|---|---|---|---|---|
| S225L | 1 (LOF) | 1 (ND) | 0 (0) | 1 (ND) | 1 (GOF) | 0 (0) | 1 (GOF) |
| S225W | 1 (UNC) | 1 (GOF) | 1 (0) | 1 (ND) | 1 (NM) | 0 (NM) | 1 (NM) |
| R228M | 1 (NM) | 1 (NM) | 0 (NM) | 1 (NM) | 1 (GOF) | 1 (GOF) | 1 (GOF) |
| Q234L | 1 (LOF) | 1 (ND) | 1 (GOF) | 1 (UNC) | 0 (NM) | 0 (NM) | 1 (NM) |
| Q234P | 1 (LOF) | 1 (ND) | 1 (LOF) | 1 (ND) | 0 (0) | 0 (0) | 1 (0) |
| Q234R | 1 (LOF) | 1 (ND) | 1 (UNC) | 0 (0) | 1 (NM) | 1 (NM) | 1 (NM) |
| R237M | 1 (LOF) | 1 (GOF) | 1 (UNC) | 1 (UNC) | 0 (0) | 0 (0) | 1 (GOF) |
| G245R | 1 (LOF) | 1 (ND) | 1 (0) | 1 (ND) | 1 (NM) | 1 (NM) | 1 (NM) |

Predicted class: Normal=0, Dysfunctional=1

**Figure S5. Individual function and trafficking predictions for S4 set variants.** These are the class predictions used to calculate the global scores in **Figures 7C** and **7D**. (**A**) Heatmap where variants predicted as dysfunctional are marked by blue cells. Text annotations include the predicted class (normal=0, dysfunctional=1) and the true label in parentheses. LOF, GOF, and ND variants were considered "dysfunctional" for classification labels. Black text in the cells indicates the assigned class label and prediction agree. Red text indicates the prediction does not match the label. Gray text indicates that the experimental value (true label) for the variant was either uncertain ("UNC") or not measured ("NM"). The variants with codon changes are marked. In the top panel, the true label is based on the measurement using the G>C codon to express the variant. In the bottom panel, the true label is based on the measurement using the specified codon. Note the RF classifiers do not consider codon changes, therefore the predicted class of the variant remains the same. (**B**) Variants in the S4 set that were part of the model training data. These replicate measurements were collected with minor changes to the experimental protocol (see Methods) and therefore may have a different assigned label. Predicted class remains the same.

**Figure S6**

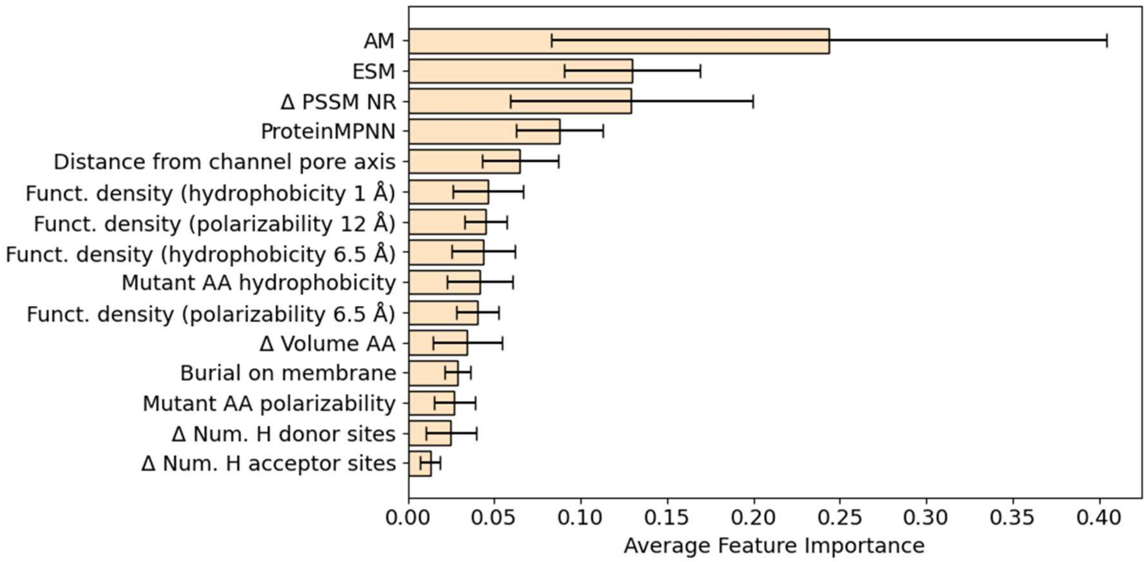


**Figure S6. RF classifier feature importance.** Average feature importance for the 15 features included in the BioEvo+AM+MPNN+ESM feature set. Feature importance values range from 0 to 1, with higher values indicating stronger contributions to model predictions. Importances were determined for each of the seven RF classifiers trained to predict protein fitness metrics. Bars and lines represent the mean and standard deviation across classifiers.

### Supplemental Tables

**Table S1**

| Variant | Surface expression (%WT) | Total expression (%WT) | Trafficking efficiency (%WT) |
|---|---|---|---|
| R228C | 347 ± 25 | 213 ± 15 | 163 ± 8.6 |
| R228M | 357 ± 28 | 210 ± 18 | 171 ± 9.0 |
| V241I | 100 ± 7.8 | 108 ± 8.6 | 94 ± 13 |

**Table S1. Trafficking measurements.** Table reports mean±STD, where N=4 for both variants. These variants were measured as part of [4] with the same experimental protocol. Both variants are considered GOF trafficking.

**Table S2**

| Measurement | Trees | Max depth |
|---|---|---|
| $I_{peak}$ | 1024 | 5 |
| $V_{1/2}$ | 128 | 2 |
| $\tau_{act}$ | 512 | 2 |
| $\tau_{deact}$ | 256 | 7 |
| Surface expression | 1024 | 5 |
| Total expression | 256 | 7 |
| Trafficking efficiency | 512 | 5 |

**Table S2. RF classifier hyperparameters for the BioEvo+AM+ProteinMPPN+ESM feature set**. For each measurement, the hyperparameters of the corresponding RF classifier were tuned independently using grid search in the Python scikit-learn library. Trees (n_estimators) are the number of trees in the forest and max depth (max_depth) is the maximum depth of each tree. These values, and those of the models using the other feature sets, are accessible in the file “saved_models/out_rfc._hyperparams.joblib”.
